# Optimizing photoautotrophic production of soluble 3-hydroxybutyrate in *Synechocystis* sp. PCC 6803 through combinatorial translational tuning

**DOI:** 10.64898/2026.01.09.698553

**Authors:** L. Kakko, D. Muth-Pawlak, P. Patrikainen, E-M. Aro, P. Kallio

## Abstract

While photosynthetic cyanobacteria are potential biotechnological hosts for light-driven production of sustainable chemicals from CO_2_, engineering more efficient strains is critical for the development of competitive industrial processes. This study demonstrates significantly enhanced production of the soluble bioplastic precursor (R)-3-hydroxybutyrate (3HB) that has been engineered based on the polyhydroxybutyrate (PHB) pathway in photoautotrophic cyanobacterium *Synechocystis* sp PCC 6803. As the key novelty, we generated a library of engineered 3HB pathway variants that express the three key heterologous pathway enzymes PhaA, PhaB and TesB at varying efficiencies, followed by the screening of most efficient 3HB producers. This was achieved by placing each of the pathway enzymes under the translational regulation of three alternative RBSs in different combinations, resulting in strains with wide dynamic range of 3HB productivities. The best strains accumulated over 5 gl^−1^ under 200 μmol photons m^−2^s^−1^ and 3% CO_2_ in a 14-day flask batch culture, with the highest titer reaching 12 gl^−1^, corresponding to nearly 3 gl^−1^ d^−1^ during the peak production phase. These are the highest 3HB production levels reported so far in cyanobacteria, and comparable to those previously established in heterotrophic production systems. Proteomic comparison of selected strains revealed that the different RBS combinations result in varying expression patterns of the pathway proteins, and that the strain-specific enzyme levels remained relatively constant over the monitored six-day period. The results show that altering the levels of the target pathway enzymes can dramatically improve product yield in *Synechocystis*, while even very small quantitative differences in the strain-specific expression profiles can have marked effects on the production efficiency. This could be used as a general tool for optimizing engineered pathways in cyanobacteria, provided that the flux to the end-product is not critically restricted by substrate availability but rather determined by the balance between the consecutive pathway steps.

## Introduction

In parallel to global electrification and the development of new bioproduction strategies, large-scale industrial biotechnologies have an increasing role in the sustainable replacement of oil-based products now in use. In this context, photosynthetic cyanobacteria have been recognized as potential next-generation biotechnological hosts that can be used for the light-driven production of carbon-based chemicals directly from CO_2_ [1]. While different cyanobacterial species have been genetically engineered to produce a wide array of industrially relevant chemicals such as butanol [2], sucrose [3], and ethylene [4], polyhydroxyalkanoates (PHAs) have gained increasing interest as a means of producing biodegradable bioplastics that can be used as alternative for petrochemical products. PHAs are biopolymers composed of small organic subunits and have various applications depending on their monomer composition, which determines their physical properties. One of the most common PHAs is polyhydroxybutyrate (PHB) composed of 3-hydroxybutyrate (3HB) monomers, which is produced naturally by various microbes. Although the PHB production levels in cyanobacteria are relatively low compared to many heterotrophic bacteria, the photoautotrophic system is an interesting research target [see reviews [5] [6]] as production does not rely on organic substrate feedstock as in heterotrophic systems, or on hydrogen as in gas fermentation. As substrate availability is a major limitation in large-scale industrial manufacturing, cyanobacterial platforms are being further developed to increase productivity and reduce costs in order to make these technologies commercially competitive in global market.

The majority of studied cyanobacteria including *Synechocystis* sp PCC 6803 (*Synechocystis* from herein) accumulate PHB as part of the normal autotrophic metabolism [7] forming compact PHB granules that may naturally constitute up to 20-25% of the cell dry weight. PHB production is primarily triggered by the limitation of the macronutrients nitrogen and phosphorous, and enhanced by day-night growth cycles [8] and supplementation of organic substrates [9] [10] and bicarbonate [11]. The process is closely linked to the breakdown of the primary carbohydrate storage polymer glycogen, which takes place mainly through the EMP pathway and provides the acetyl-CoA precursors for PHB biosynthesis under nutrient-limitation [12]. The *pha* pathway consists of three consecutive steps catalyzed by acetyacyl-CoA β-ketothiolase PhaA, acetoacetyl-CoA reductase PhaB and heteromultimeric PHA synthase PhaEC (**Figure 1**) that convert acetyl-CoA to PHB. The production efficiency of PHB has been enhanced by overexpressing the *pha* proteins in *Synechocystis* [13] [14] [15] and in *Synechococcus elongatus* UTEX 2973 [16], by improving the carbon capture by overexpressing RuBisCo [17], and modification of proline and arginine metabolism to increase the flux to PHB via acetyl -CoA under nutrient limitation [18]. Strains have also been engineered by the deletion of PirC, central carbon flow regulator that responds to nitrogen availability, significantly increasing carbon flux towards PHB under nutrient deprivation [10] and by the inactivation of the phosphate regulator SphU that results in the stimulation of PHB biosynthesis even under nitrogen-sufficient conditions [19]. By combining conditional optimization of the cultures through nutrient limitation, supplementation of organic substrates and optimization of the light- and carbon conditions, the highest PHB levels have reached over 80% of the dry cell [10]. Despite the advancements, industrial production would also require more efficient means of cell harvesting, large scale cell disruption and PHB recovery, all of which add up to the production costs [6].

**Figure 1.**
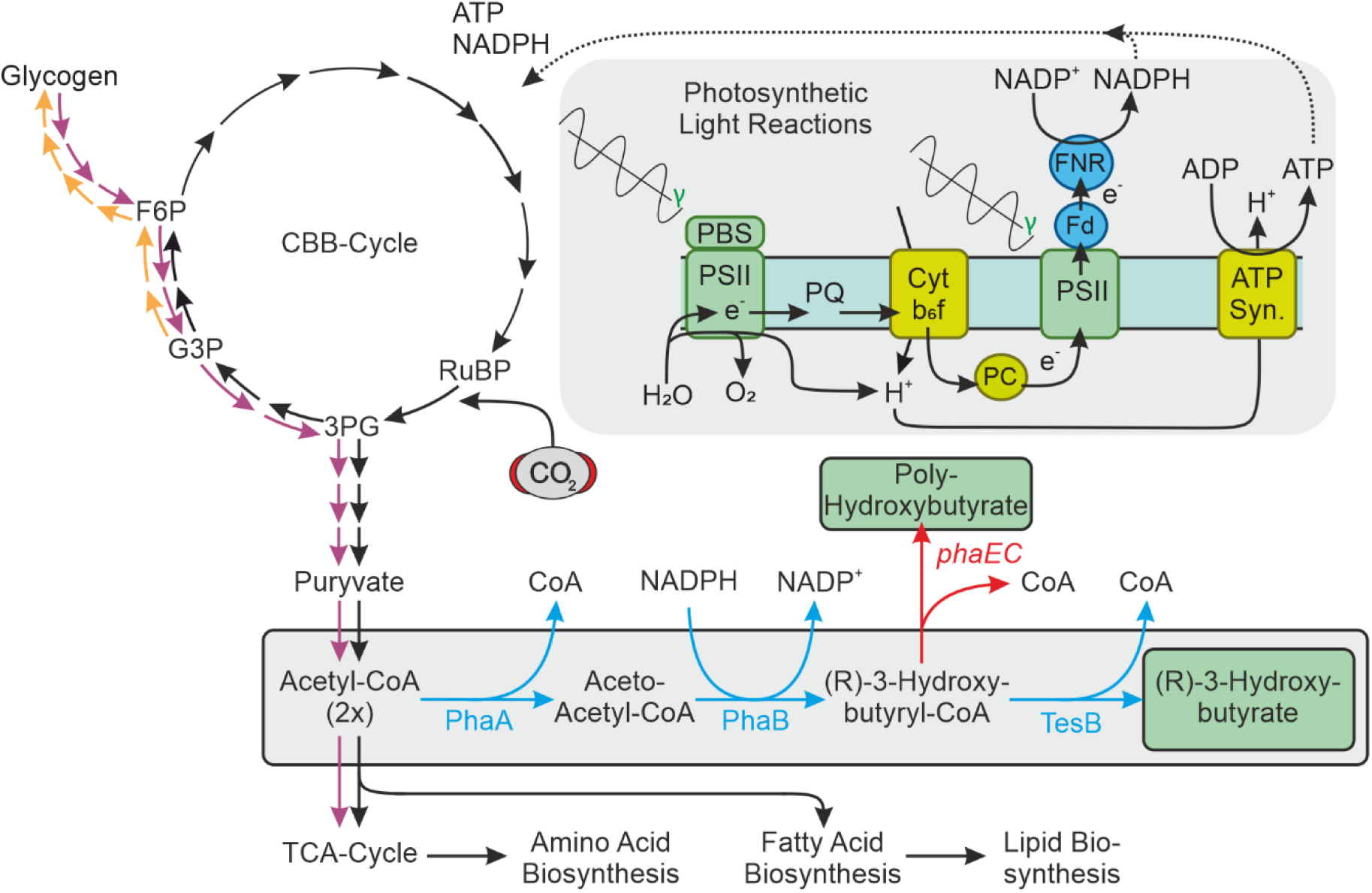
Representation of the pathway engineered for the production of 3HB in *Synechocystis*. The engineered 3HB pathway is composed of consecutive enzymatic reactions catalyzed by three overexpressed heterologous enzymes PhaA, PhaB and TesB (blue arrows) based on the original design of [28]. PhaA is a PHA-specific β-ketothiolase (*Ralstonia eutropha* H16; EC:2.3.1.9) that condensates two acetyl-CoA units to form acetoacetyl-CoA. PhaB is a PHA-specific acetoacetyl-CoA reductase (*Ralstonia eutropha* H16; EC:1.1.1.36) that reduces acetoacetyl-CoA into 3-hydroxybutyryl-CoA using NADPH as cosubstrate. TesB is a Acyl-CoA thioesterase 2 (*Escherichia coli*; EC:3.1.2.20) that cleaves off CoA from 3-hydroxybutyryl-CoA, releasing monomeric 3HB that secretes out from the cell. The native PHB biosynthesis is blocked by the deletion of *phaE* and *phaC* encoding the PHB synthase PhaEC. 3HB production is powered by the photosynthetic machinery of the *Synechocyctis* host. The cell uses NADPH and ATP generated in the photosynthetic light reactions to fix CO_2_ into G3P that is the initial precursor for various metabolic reactions. Large fraction of the flux to the 3HB pathway proceeds via glycogen, which is produced as a storage compound (yellow arrows) and broken down through the Embden-Meyerhof-Parnas (EMP) pathway to acetyl-CoA (purple arrows) under the production conditions. Abbreviations: ATP, Adenosine triphosphate; CBB-Cycle, Calvin–Benson–Bassham cycle; CoA, Coenzyme A; Cytb_6_f, Cytochrome b6f; Fd, Ferredoxin; FNR, Ferredoxin-NADP^+^ reductase; F6P, Fructose-6-phosphate; G3P, Glyceraldehyde-3-phosphate; 3PG, 3-phosphoglycerate; PBS, Phycobilisome; PSI, PC, Plastocyanin; NADP^+^/NADPH Nicotinamide adenine dinucleotide phosphate; Photosystem I; PSII, Photosystem II; PQ, Plastoquinone; RuBisCO, ribulose-1,5-bisphosphate carboxylase-oxygenase; RuBP, Ribulose 1,5-bisphosphate; TCA-Cycle, Tricarboxylic acid cycle; γ, photons in the 400–700 nm PAR range.

Studies suggest that the efficient over-production of PHB in cyanobacteria is challenging due to slow accumulation, direct correlation with the biomass content and difficulty in bypassing the native regulatory circuits. One strategy to overcome these constraints is to prevent the native PHB polymer formation and engineer the cells to excrete monomeric 3HB that accumulates outside the cell in the medium. This may possibly circumvent endogenous limitations caused by end-product inhibition, and enable continuous bioproduction strategies that do not require host cell lysis to harvest the product. This has been implemented in several heterotrophic microbes including *Escherichia coli* [20] [21] [22] and methanotrophic bacteria *Methylotuvimicrobium alcaliphilum* [23], yeasts *Saccharomyces cerevisiae* [24] and *Arxula adeninivorans* [25] as well autotrophic acetogenic bacteria such as *Clostridium ljungdahlii* that utilize H_2_ and CO_2_ for the production of 3HB [26] [27]. To further improve the sustainability of the process, several studies have focused on the light-driven photoautotrophic production of 3HB from CO_2_ by genetically engineering cyanobacteria [28] [29] [30]. This strategy is generally based on i) inactivating the native PHB biosynthesis by deleting the PHB synthase genes *phaEC*, ii) enhancing the conversion of acetyl-CoA to 3-hydroxybutyryl-CoA via acetoacetyl-CoA, and iii) detaching the CoA moiety by expressing a heterologous thioesterase to allow 3HB excretion from the cell.

As demonstrated in the engineered 3HB pathways in cyanobacteria, one of the key challenges in strain development is to balance the consecutive enzyme-catalysed conversion steps in artificial biosynthetic pathways to achieve maximal flux to the end product [29] [30]. Imbalance between the reactions may lead to intermediate accumulation and harmful physiological effects that negatively affect the host and reduce productivity [30]. Our objective here was to address this issue further, and evaluate the potential of translational pathway tuning [31] for enhancing the 3HB production in genetically engineered *Synechocystis*. The strategy was to use a combinatorial approach to balance the key steps in 3HB biosynthesis in concert, by expressing the three enzymes PhaA, PhaB and TesB at different relative efficiencies, followed by screening of the highest producing strain variants with the most optimum combination. This can be accomplished by using alternative RBS-elements upstream each pathway gene to control target protein translation [32], thus enabling the generation of a library of strains with varying expression profiles. Although translational tuning has been shown to be effective in optimizing multi-step pathways in *E. coli* [31], it has only been implemented in a few cases in cyanobacterial engineering [33] [34] [35]. In addition to improving the productivity of 3HB, the goal was to assess the potential of translational tuning as a general engineering tool for developing enhanced photoautotrophic strains for future use in biotechnology.

## Materials & Methods

### Chemicals, reagents and DNA synthesis

All enzymes including DNA polymerases Taq and Phusion, T4 DNA ligase and the restriction enzymes were purchased from New England BioLabs (US). Commercial kits including QIAprep Spin Miniprep Kit, CompactPrep Plasmid Midi Kit and QIAquick Gel Extraction Kit were purchased from Qiagen (DE). PCR primers were ordered from Eurofins MWG Operon (DE) and the synthetic genes from GenScript (US). Rest of the reagents and chemicals were purchased from Sigma-Aldrich (US), unless stated otherwise.

### Microbial strains and cultivation conditions

*Escherichia coli* strain DH5α was used for plasmid amplification and construct library generation in this work. The cells were routinely grown at 37 °C in Luria Bertani (LB) liquid medium in 250 rpm shaking or LB agar plates (1% w/v). Antibiotics were supplemented at maximum concentrations 100 µgml^−1^ ampicillin (pNiv selection), 50 µgml^−1^ kanamycin (pSI1NL-KmR selection), 50 µgml^−1^ spectinomycin (pDF selection), 34 µgml^−1^chloramphenicol (gene insert selection) unless specified otherwise.

The sucrose-tolerant cyanobacterium *Synechocystis* sp. PCC 6803 (Kaplan) was used as the background strain for engineering. All cultures were grown in BG11 liquid medium pH 8.3 buffered with TES-KOH in either 10ml volume in 25 ml Erlenmeyer flasks (precultures) or in 20ml volume in 50ml Erlenmeyer flasks (analytical cultures), or on corresponding agar plates. Antibiotics were supplemented at maximum concentrations 25 µgml^−1^ spectinomycin (pDF selection), 50 µgml^−1^ kanamycin (Δ*phaEC* selection) and 10 µgml^−1^chloramphenicol (insert selection) unless specified otherwise. The precultures and plates were grown in Sanyo MLR-351 environmental test chamber under constant white light illumination 50 μmol photons m−^2^ s−^1^ (1% CO_2_) at 30 °C with 120 rpm shaking. The main cultures were grown in a POL-EKO Apartura culture cabinet using a customized light system to ensure uniform illumination **(Supplementary Figure S1)** under 50 μmol photons m^−2^s^−1^ (1% CO_2_) or 200 μmol photons m^−2^s^−1^ (3% CO_2_) at 30°C in 120 rpm shaking. 3HB production was induced by supplementing isopropyl β-D-1-thiogalactopyranoside (IPTG) at the beginning of the main culture (OD_750nm_ = 0.5) in 1mM final concentration.

### Assembly of the construct library in *E. coli*

Assembly of the expression plasmids followed the original cloning protocol used in [31] that we have later adapted for *Synechocystis* [32]. The genes *tesB*, *phaB* and *phaA* were purchased as separate synthetic DNA fragments fused to chloramphenicol resistance cassette (CmR) with flanking restriction sites *Nsi*I-*Xho*I and *Nhe*I in between the gene and CmR (**Supplementary Figure S2**). Each gene was first combined with three alternative RBS sequences (S3: AGTCAAGTAGGAGATTAATTCAATG, S4: ATACATAAGGAATTATAACCAAATG, B: AGGAGGTTTGGAATG) [32] by subcloning the ordered fragment using *Nsi*I-*Xho*I into a mixture of corresponding pNiv assembly plasmids. In the second step the target genes with the RBSs were fused together in two iterative rounds of subcloning using the compatible enzyme pairs NheI/SpeI XhoI/SalI in pNiv as described earlier [31]. The resulting mixture of constructs carrying the three target genes combined with three alternative RBSs were subcloned into the pDF-lac2 expression vector backbone using SpeI-SalI to generate the library of pDF-lac2-(x)tesB-(x)phaB-(x)phaA-CmR constructs.

After each transformation step the *E. coli* clone library was grown on LB agar plates containing either 100 µgml^−1^ ampicillin and 34 µgml^−1^ chloramphenicol (selecting for pNiv assembly plasmids with the insert) or 50 µgml^−1^ spectinomycin and 34 µgml^−1^ chloramphenicol (selecting for pDF expression plasmids with the insert). The plasmids were extracted by adding 1-3ml of LB medium onto the plates, suspending the colonies in the liquid by scraping with a soft plastic loop, and promptly extracting the plasmids from the resulting cell suspension using Qiagen plasmid Miniprep or Midiprep kit. To ensure that that all the constructs were well represented in the library throughout the procedure, each extraction typically combined at least 500 *E. coli* colonies, sometimes from multiple parallel transformant plates.

### Generation of the *Synechocystis* Δ*phaEC* strain

To assemble the vector for deleting *phaEC* by homologous recombination, the primer pairs phaE(XbaI)_F/phaE(XmaI)_R and phaC(AvrII)_F/phaC(ClaI)_R were used for amplifying ∼500 bp fragments from the beginning of *phaE* and the end of *phaC* by PCR using *Synechocystis* genomic DNA as template. The amplified sequences were inserted on the both sides of kanamycin resistance cassette (KmR) in pSI1NL-KmR using the restriction enzyme pairs XbaI/XmaI and ArvII/ClaI to generate the final construct pSI1NL-KmR-phaEC. The pSI1NL-KmR used as the backbone is a derivative of pSI1B [36] where the region spanning P_A1lacO-1_ and LacI has been replaced by KmR. To delete *phaEC*, pSI1NL-KmR-phaEC was amplified in *E. coli* and transformed in *Synechocystis* WT using natural competence [37]. First a 5 ml aliquot of liquid cell culture at the mid logarithmic growth phase (OD_750nm_ ∼0.8) was gently pelleted and resuspended in 1 ml fresh BG11. The plasmid was added in the suspension to the final concentration 20 µg/ml, and the mixture was incubated for 5h at 30 °C, 1% CO_2_ with 120 rpm shaking under constant illumination 50 μmol photons m−^2^ s−^1^. The cells were then transferred on a fresh BG11 plate for two days, followed by supplementation of kanamycin (10 µgml^−1^) underneath the agar. After the appearance of colonies, the transformant cells were streaked on secondary plates with increasing concentrations of kanamycin (up to 50 µgml^−1^). The presence of the KmR insert and segregation of *ΔphaEC* were confirmed by colony PCR using the primer pairs phaEC_F/ KmR_R and phaEC_F/phaEC_R (**Table 2**), respectively.

**Table 1:**
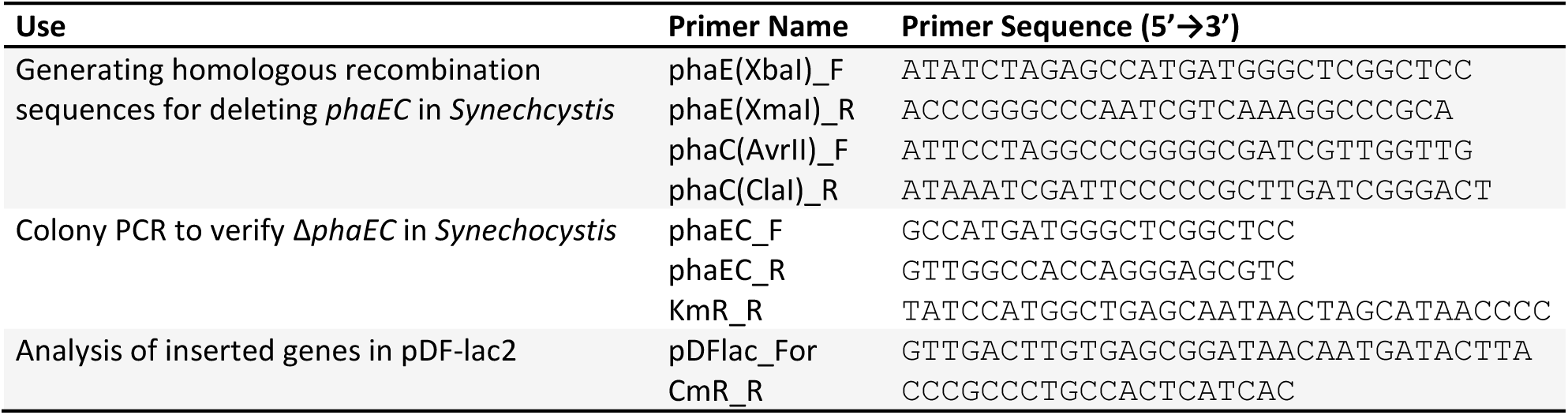
The PCR primers used in this study.

**Table 2.** Cell strains and plasmid constructs generated and used in this study. (x) refers to randomized RBS generated by using a mixture of the three RBSs.

| Strain | Description | Source |
| --- | --- | --- |
| *E. coli* DH5α | *E. coli* strain used for plasmid amplification and cloning | Invitrogen |
| *Synechocystis* WT | The background sucrose-tolerant strain "*Kaplan*" | Aaron Kaplan |
| *Synechocystis* Δ*phaEC* | The 3HB library background strain; inactive PHB synthesis | This study |
| *Synechocystis* Δ*phaEC* 3HB strains 001-273 | Library of *Synechocystis* Δ*phaEC* strains w/ the pDF-lac2-(x)tesB-(x)phaB-(x)phaA-CmR construct variants | This study |
| **Plasmid** |  |  |
| pUC57-Simple-tesB-CmR | Synthesized *tesB* and CmR-cassette in carrier plasmid | This study |
| pUC57-Simple-phaA-CmR | Synthesized *phaA* and CmR-cassette in carrier plasmid | This study |
| pUC57-Simple-phaB-CmR | Synthesized *phaB* and CmR-cassette in carrier plasmid | This study |
| pNiv(S3) | pNiv assembly plasmid carrying the RBS S3 | [32] |
| pNiv(S4) | pNiv assembly plasmid carrying the RBS S4 | [32] |
| pNiv(B) | pNiv assembly plasmid carrying the RBS B | [32] |
| pNiv(x)tesB-CmR | pNiv library; tesB-CmR w/ RBS S3, S4 and B | This study |
| pNiv-(x)phaA-CmR | pNiv library; phaA-CmR w/ RBS S3, S4 and B | This study |
| pNiv-(x)phaB-CmR | pNiv library; phaB-CmR w/ RBS S3, S4 and B | This study |
| pNiv-(x)tesB-(x)phaB-CmR | pNiv library; tesB-phaB-CmR w/ randomized RBSs | This study |
| pNiv-(x)tesB-(x)phaB-(x)phaA-CmR | pNiv library; tesB-phaB-phaA-CmR w/ randomized RBS | This study |
| pDF-lac2-(x)tesB-(x)phaB-(x)phaA-CmR | pDF-lac2 library; tesB-phaB-phaA-CmR w/ randomized RBSs | This study |
| pSI1B | Plasmid backbone for *phaEC* deletion vector assembly | [36] |
| pSI1NL-KmR | Incomplete deletion vector carrying KmR | This study |
| pSI1NL-KmR-phaEC | *phaEC* deletion vector w/ recombination sequences and KmR | This study |

### *Synechocystis* library transformation and maintenance

The pDF expression plasmid library extracted from *E. coli* was transformed in repeated cycles in *Synechocystis* Δ*phaEC* using the same procedure as described above. For transformant selection, the cells were first transferred on fresh BG11 agar plates for 2-4 days, after which antibiotics were administered underneath the agar to 20% (5 µg/ml spectinomycin, 10 µg/ml kanamycin and 2 µg/ml chloramphenicol) of the final concentration. Colonies that typically appeared on the plates within 10-30 days were transferred to secondary plates containing 50%, and finally the full amount of antibiotics (25 µg/ml spectinomycin, 50 µg/ml kanamycin and 10 µg/ml chloramphenicol). Each independent colony was numbered and transferred on a gridded square BG11 agar plate with antibiotics, and the process was repeated until 273 transformants had been generated (**Supplementary Figure S3**). To maintain the library, the colonies were transferred on fresh plates at six-week intervals using a 3D-printed autoclavable stamp tool designed for the grid. The new plates were incubated for one week at 30 °C, 1% CO_2_, 50 μmol photons m−^2^ s−^1^, after which they were stored on a lab shelf at ambient temperature and light for 5 weeks.

### 3HB sampling and derivatization

Analytical samples for 3HB quantitation (V = 0.5 ml) were collected from the liquid cultures at the given time points and stored at -80°C until use. The samples for GC-MS analysis were first dried under ambient air in 10µl sample aliquots in glass GC-MS vial with a glass insert (Crimp vial with fixed insert, Agilent, DE) for 16 hours in 42°C. To derivatize 3HB, the evaporated samples were resuspended to 25µL of nonane, and derivatized by silylation by adding 25µL BSTFA+1%TMCS for 4h at 70°C [38]. Each prepared sample set was subjected to GC-MS analysis immediately after derivatization.

### Preparation of the 3HB standard curve

The quantitation of 3HB in the culture samples was based on comparison with commercial 3HB standard (95% 3-hydroxybutyric acid, Sigma-Aldrich, US). The standard dilutions were prepared by first weighing and carefully solubilizing the commercial 3HB stock to the final concentration of 25 mgml^−1^ in 10 ml BG11, followed by independent dilutions to concentrations 1, 5, 10 and 15 gl^−1^ in three parallels. After careful mixing, aliquots of the standard solutions were subjected to sample treatment and derivatization exactly as described for the culture samples. After CG-MS analysis the obtained 147 m/z signals corresponding to 3HB were plotted against the standard concentrations (**Supplementary Figure S4**), and used determining the concentrations of the culture samples over the linear range between 1-15 gl^−1^.

### GC-MS analysis and 3HB quantitation

GC-MS analysis of 3HB [38] was performed using Agilent 7890A gas chromatography instrumentation equipped with an autosampler and 5975C Inert MSD with Triple Axis Detector (Agilent), and DB-5MS ultra inert fused silica capillary column 30m (Agilent). The derivatized samples were injected in 1µl volume in 1:200 split mode (He carrier gas flow 1 mlmin^−1^, inlet temperature 250 °C) followed by temperature gradient run (initial oven temperature of 50 °C for 1 min, increased by 20 °C min^−1^ to 150 °C, held for 1 min, and increase by 35 °C min^−1^ to 320 °C). The 147 m/z peak corresponding to 3HB eluted at around 595 s and quantification was done based on the manually integrated peak area and the 3HB standard curve.

### Sample preparation for proteomic analysis

The samples (V_tot_ = 2.5 ml) for the quantitative proteomic analysis of TesB, PhaB and PhaA were collected at the three given time points (d8, d11, d14) from three replicate liquid cultures of the seven selected strains, pelleted and stored at -80°C until use. The proteins were isolated and digested using the protocol described previously [39] [40] with small modifications. The cell pellets were suspended in 150 µl buffer containing 6M urea in 0,1M Tris-NaOH pH 8 buffer, 1% PMSF, mixed with an equal volume of glass beads and lysed using a bead beater. A volume of the crude lysate corresponding to 100 µg of total protein was then subjected to reduction with DTT, and alkylation with IAA, followed by precipitation of proteins in cold acetone/ethanol mixture in -20°C overnight. The proteins were then digested with trypsin in 0,05 M Tris-NaOH pH 8 buffer for 20 h. The mixture of peptides was desalted via Sep-Pack C18 (Waters) columns with the protocol recommended by the manufacturer. Samples were diluted to ensure 500 ng of peptides injection and spiked with a mixture of 10 heavy peptides (PepoTech 2^nd^ Grade, Thermo Scientific) to reach around 100 fmolµl^−1^ concentration reference to quantify three engineered proteins (P0AGG2, P14697 and P14611) (**Supplementary Table S1**).

### Proteomic analysis of the 3HB pathway proteins

The LC-ESI-MS/MS analysis was performed for all the 64 samples on a nanoflow HPLC system (Easy-nLC 2000, Thermo Fisher Scientific) coupled to the QExactive-Orbitrap mass spectrometer (Thermo Fisher Scientific) equipped with a nano-electrospray ionization source. The injected samples were first trapped on pre-column and then separated on an analytical C18 column (75 μm x 15 cm, ReproSil-Pur 3 μm 120 Å C18-AQ, Dr. Maisch HPLC GmbH, Ammerbuch-Entringen, DE) by a two-step, 60 min gradient from 5 to 26% solvent B over 35 min, followed by 26 to 49% B increase over 15 min. The mobile phase consisted of water with 0.1% formic acid (solvent A) or acetonitrile/water (80:20 (v/v)) with 0.1% formic acid (solvent B).

The MS data was acquired automatically by using Thermo Xcalibur 4.1 software (Thermo Fisher Scientific). A Parallel Reaction Monitoring (PRM) method was applied to monitor 10 peptides representing the three target proteins. The acquisition method consisted of an Orbitrap MS survey scan of mass range 300-2000 m/z followed by a set of targeted MS^2^ (tMS^2^) HCD scans to fragment the ions according to scheduled inclusion list (**Supplementary Table S1**). The spectra were registered with a resolution of 120 000 and 60 000 (at m/z 200) for MS and tMS^2^, respectively, and normalized collision energy of 27%. The automatic gain control (AGC) was set to a maximum fill time of 200 ms and maximum number of 3e6 ions for MS scan while for tMS^2^ maximum fill time was set to 110ms or 1e6 ions.

The generated raw files were deconvoluted to recover peptide fragments from PRM data in Skyline [41] software. The sequence of the peptides was confirmed with reference spectra from a spectral library built in-house. The chromatograms of peptide fragments representing heavy and light peptides were manually checked for correct integration. The light/heavy ratios for peptides were exported and further processed in Excel.

### Statistical analysis of the proteomic data

To evaluate the statistical significance in the pathway protein abundances between the strains, the highest producer 33B was used as the reference for the comparison. The statistical analysis of the protein abundances was performed by pooling together all the validated proteomic data from the three sampling points for each strain. As the data exhibited widespread non-normal distribution (confirmed by Shapiro-Wilk tests, with p < 0.05 for 20 of 21 datasets) and unequal sample sizes, the Mann-Whitney U-test was used for evaluating the significance between the expression level differences. An approximate z-test was used to assess differences in protein ratios calculated on the log scale, as it effectively compares derived ratios from non-normally distributed abundance data (log-transformed to stabilize variance). The method was selected for its suitability moderately large sample sizes (N ≥ 18), where the Central Limit Theorem justifies the approximate normality of mean differences, and it accommodates unequal sample sizes and variances. To evaluate the differences in the protein ratios TesB:PhaA, TesB:phaB and PhaA:PhaB, the overall ratio was calculated as the arithmetic mean of the numerator protein divided by the arithmetic mean of the denominator protein. Cohen’s d was used measure of effect size quantifying the standardized difference between the means of log-ratios of TesB:PhaA, TesB:phaB and PhaA:PhaB pairs between the reference strain 33B and other 6 comparator strains.

## Results

### Design of the strategy for optimizing 3HB production in *Synechocystis*

The platform for optimizing photoautoprophic 3HB production in *Synechocystis* was based on modified endogenous PHB pathway, following an engineering strategy presented earlier [28] [29]. The adapted biosynthetic route utilizes the intracellular acetyl-CoA as the precursor, and relies on the overexpression of three heterologous biosynthetic enzymes, PhaA (acetyacyl-CoA β-ketothiolase; *Ralstonia eutropha* H16), PhaB (NADPH-dependent acetoacyl-CoA reductase; *Ralstonia eutropha* H16) and TesB (acyl-CoA tioesterase; *E. coli*) (**Figure 1**) [28]. When the native PHB polymerization reaction is blocked in the background strain by the deletion Δ*phaEC*, the three overexpressed enzymes convert available acetyl-CoA in consecutive reaction steps into a monomeric 3HB, which is excreted from the *Synechocystis* cell and accumulates in the culture medium [28]. With the objective of applying translational tuning for optimizing the relative efficiency of the consecutive conversion steps towards 3HB, the first goal was to design pathway variants with different translational patterns between the target proteins PhaA, PhaB and TesB. To regulate the expression, three alternative translational control elements were selected from an RBS library previously characterized in *Synechocystis* [32]. These specific RBSs, S3 (derived from *cpcB sll1577* in *Synechocystis*), S4 (derived from *psbA2 slr1131* in *Synechocystis*) and B (reverse engineered for *E. coli*) [42] were expected to provide sufficient variation in the target protein expression levels, while maintaining the library appropriately small for convenient handling.

### Generation of the Δ*phaAC* background strain for 3HB production

In order to prevent the native polymerization of 3-hydroxybutyryl-CoA into PHB in the 3HB production strain, the consecutive genes encoding for the heterooligomeric PHB synthase PhaEC (*phaE* / *slr1829* and *phaC / slr1830*) were inactivated by the insertion of a kanamycin resistance cassette (KmR) in WT *Synechocystis* by homologous recombination. This was accomplished by constructing a deletion vector pSI1NL-phaEC-KmR carrying homologous sequences from the beginning of *phaE* and the end of *phaC* on each side of KmR, followed by transformation in *Synechocystis*. After selection, the resulting kanamycin-resistant clones were analyzed by colony PCR to confirm *phaEC* deletion and complete segregation of *Synechocystis* Δ*phaEC* (**Figure 2**; **Table 2**).

**Figure 2:**
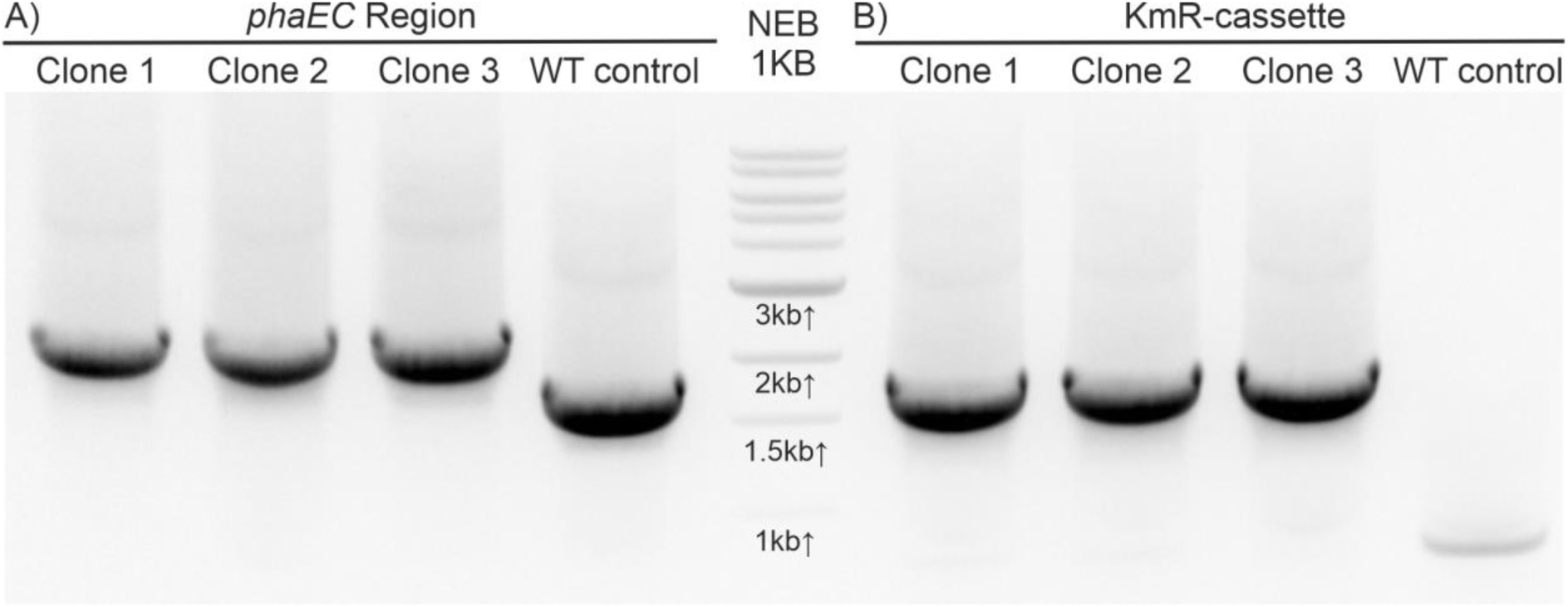
Colony PCR verification of *phaEC* deletion and segregation in the generated *Synecocystis* Δ*phaEC* mutant. **A)** PCR primers that anneal at the beginning of *phaE* and at the end of *phaC* produce a 2124 bp amplification fragment in Δ*phaEC* strain (lanes 1-3) and a 1603 bp band in WT (lane 4). B) PCR primers that anneal at the beginning of *phaE* and at the end of the inserted KmR cassette produce a 1722 bp amplification fragment in Δ*phaEC* strains (lanes 1-3) and no band in WT.

### Construction of the 3HB expression pathway variants by RBS shuffling

The 3HB pathway expression in *Synechocystis* was based on an operon composed of the three genes *phaA*, *phaB* and *tesB* arranged under the transcriptional control of an IPTG-inducible lactose promoter variant P_A1lac0-1_ in the replicative plasmid pDF-lac2 [32] [43] (**Figure 3**). The plasmid-based system was selected for the purpose since the transformants are reasonably easy to acquire as chromosomal segregation is not required, making it well applicable for obtaining a high number of clone variants required in this work. In addition, the plasmid can be conveniently amplified and selected in *E. coli* which enables relatively high expression levels in comparison to integrated chromosomal constructs and has shown to be relatively stable in *Synechocystis* [36]. The construct assembly followed an iterative subcloning strategy [32], which allowed the introduction of a desired RBSs in front of each gene, and the arrangement of the target genes as a single transcription unit in pDF-lac2. For this purpose, the three target genes *phaA*, *phaB and tesB* [28] were ordered as synthetic fragments, after omitting internal restriction sites SpeI and SalI utilized in the assembly (**Supplementary Figure S2**).

**Figure 3:**
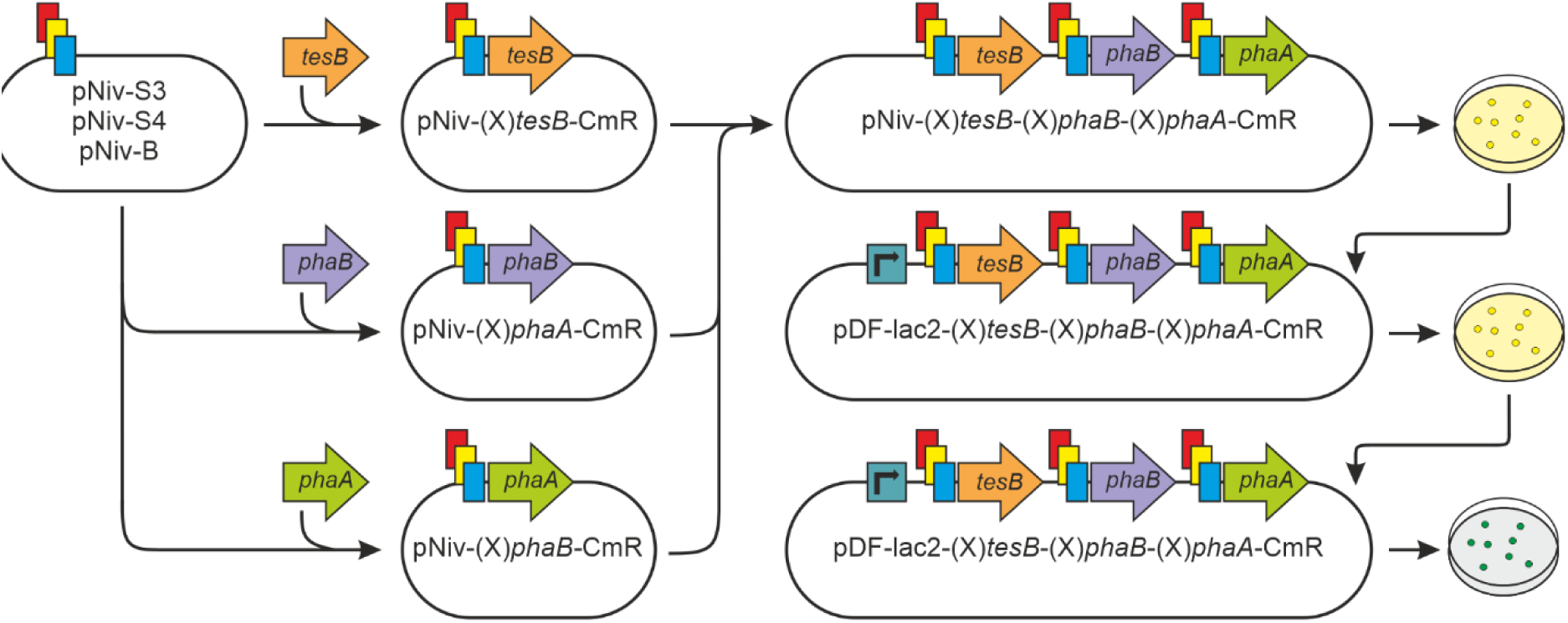
Graphical representation of the successive subcloning steps in the 3HB construct assembly. Each ordered target gene sequence (*tesB*, *phaB* and *phaA*) was first fused with the three selected RBSs in pNiv assembly vectors and then compiled together as a library of three-gene constructs. After amplification in *E. coli* the mixed library was subcloned into the pDF expression plasmid backbone, and ultimately transformed in *Synechocystis* Δ*phaEC*. The CmR selection markers used in the cloning process have not been drawn. The strategy is adapted from [31] using the RBSs described in [32].

As the proteins encoded by genes located at the beginning of an operon may be expressed more effectively [44], the genes in the 3HB operon were arranged in the specific order *tesB*, *phaB* and *phaA* to prioritize the enzymes more likely to limit the flux [29]. Maintaining the order and combining each gene with the three alternative RBSs elements (S3, S4, B) gives rise to 27 (i.e. 3^3^) distinct 3HB pathway permutations. Instead of building each of the 27 constructs one at a time, we adapted a randomized subcloning strategy for the assembly [31] (**Figure 3)** in order to study the generation and maintenance of cyanobacterial strain libraries in practice. To achieve this, the three RBSs were first fused with the target genes by using an equimolar mixture of S3, S4 and B fragments in each corresponding subcloning step. This was followed by the assembly into a library of (x)tesB-(x)phaA-(x)phaB plasmid constructs (“x” representing the randomly selected RBS element) containing all the 27 variations of the pathway. Ultimately, the three-gene fragments were subcloned into the pDF backbone to generate the expression plasmid library pDF-lac2-(x)tesB-(x)phaA-(x)phaB-CmR that was verified by PCR and restriction digestion analysis (**Figure 4**; **Table 2**).

**Figure 4:**
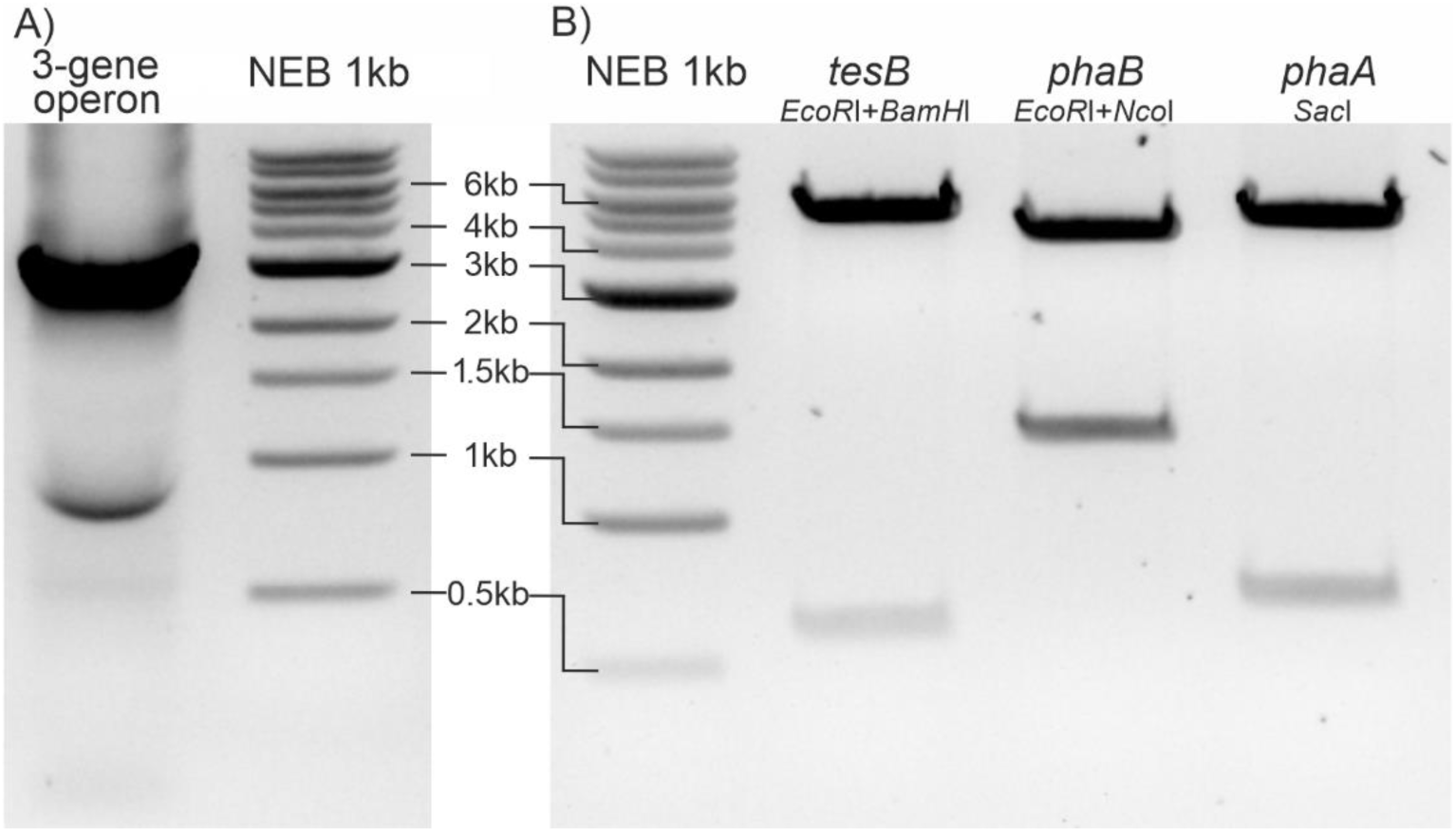
Verification of constructed expression plasmid libraries pDF-lac2-(x)tesB-(x)phaB-(x)phaA-CmR after extraction from *E. coli* prior transformation in *Synechocystis*. **A)** A representative analytical PCR reaction using primers that anneal upstream the 3HB operon in the plasmid backbone and downstream the operon in the CmR cassette to confirm the presence of all the three genes (expected size ∼3150 bp), as opposed to incomplete operons with only two genes (expected size ∼1970 bp) or a single gene (expected size ∼1230 bp). **B)** A representative restriction digestion to confirm the presence of each of the pathway genes using gene-specific enzymes *Ec*oRI and *BamH*I for *tesB* (expected sizes 5558 bp and 605 bp) *EcoR*I and *Nco*I for *phaB* (expected sizes 4690 bp and 1473 bp), and *Sac*I for *phaA* (expected sizes 5434bp and 693bp). The successive construct library assembly steps (Figure 3) were monitored accordingly.

After several successive construction rounds, the resulting plasmid mixtures were transformed in *E. coli* DH5α for amplification, giving rise to thousands of chloramphenicol / spectinomycin resistant colonies on selective LB agar plates. To minimize uneven representation of clonal variants that may result from transcriptional leakage and altered growth phenotypes in the population [45] the transformant libraries were never cultivated in liquid media. Instead, the colonies were maintained on plates until plasmid library extraction for subsequent transformation in *Synechocystis*.

### Generation and maintenance of the 3HB producer strain library

The plasmid mixtures extracted from the *E. coli* libraries were used for transforming *Synechocystis* Δ*phaEC* in more than 10 rounds of transformation and stepwise selection on antibiotic-containing BG11 plates over a course of about 20 months. Successful confirmed transformants were transferred to a library plates, which were periodically renewed and maintained at RT until over 270 independent confirmed clones had been collected. With the 27 possible RBS permutations in the expression constructs, this library covers all the different strain variations with over 98% probability (**Supplementary Figure S3**).

### Preliminary screening of the 3HB producer library to select clones for comparative analysis

Each of the 273 independent *Synechocystis* transformants were inoculated from the library plates in 10ml BG11 in 25ml conical flasks, and cultivated under 1% CO_2_ and constant white light 50 μmol photons m^−2^s^−1^ illumination for five days. To synchronize cell growth between the flasks, the precultures were used for inoculating corresponding main cultures at starting OD_750nm_ = 0.5, which were then induced and incubated under the same conditions for 14 days until 3HB quantitation. Samples were collected from each culture, derivatized and analyzed in three replicates by GC-MS for the ion m/z 147 representing 3HB (**Figure 5)**. Altogether 262 (96%) of the analyzed clones were shown to accumulate 3HB in the medium at detectable levels, while no product was observed in the WT control. Although the overall 3HB levels were low and mostly outside the linear quantitation range of the method (below ∼1mg/ml) (**Supplementary Figure S4**), the screening provided sufficient information for selecting a subset of strains from the library for subsequent analysis. The accumulation profile was clearly dynamic, and enabled us to select candidates over a relatively broad 3HB production range (**Figure 5**). Initially 30 clones were screened by sequencing to specify the RBS sequences upstream each target gene, from which 15 clone candidates were chosen for more precise quantitative characterization. These included several clones with the same RBS combinations but different 3HB production levels, indicating changes elsewhere in the construct or the host strain. The strains were named using three-digit IDs based on the order of the RBSs in the 3HB operon, with the superscripts referring to clones with different phenotypes.

**Figure 5:**
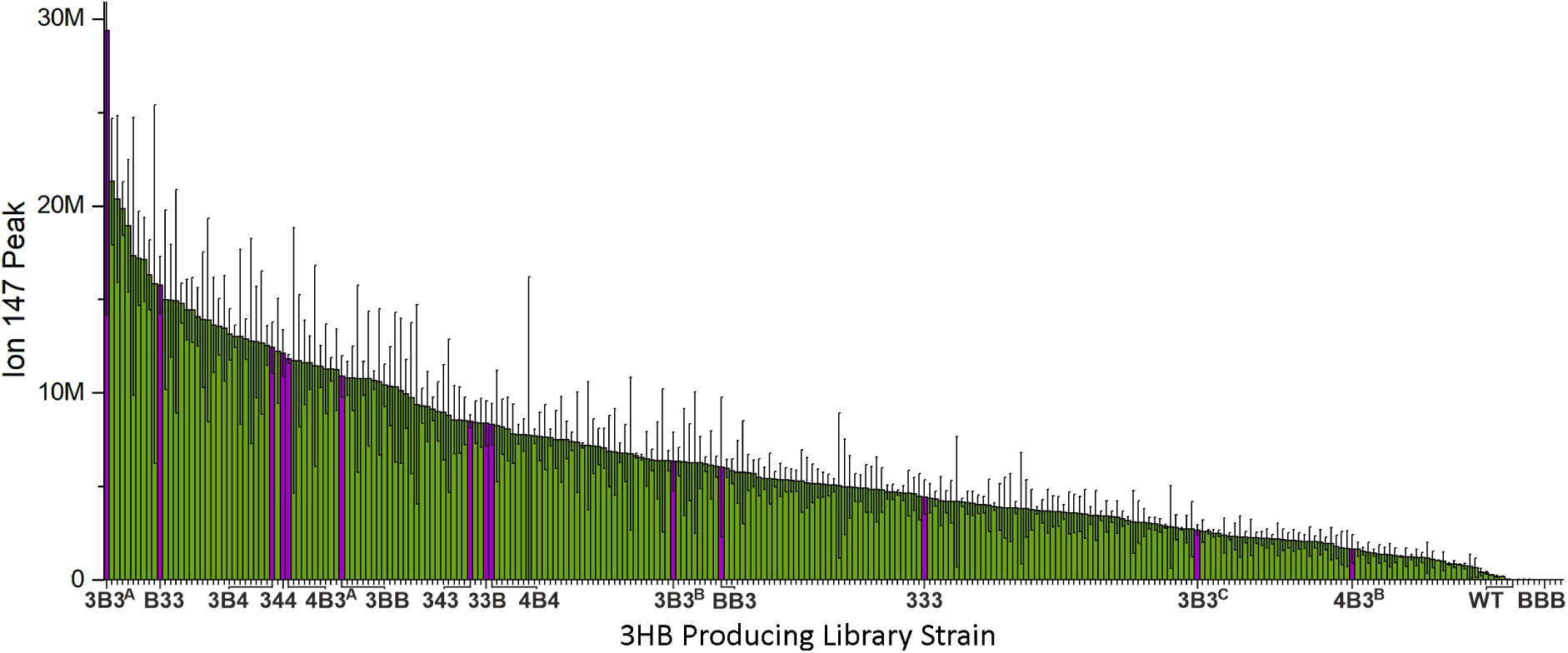
Initial screening of the generated strain library composed of 273 independent *Synechocystis* transformants engineered to produce 3HB. The strains were grown for 14 d under 50 μmol photons m−^2^ s−^1^ illumination, 1% CO_2_ in 10ml BG11 in 25ml Erlenmeyer flasks, and analyzed for 3HB. The productivities are represented as integrated m/z ion 147 peak areas that correspond to 3HB, as some of the values reside outside the linear range of the standard curve (**Supplementary Figure S4**). The averages and standard deviations have been calculated from a single culture based on three technical replicates (n = 3). The 15 strains selected for the subsequent round of analysis are shown as purple bars. The three-digit strain names refer to the order of the RBSs in the 3HB operon (e.g., “3B4” indicating (RBS S3)*tesB*-(RBS B)*phaA*-(RBS S4)*phaB*). The superscripts refer to distinct clones with the same RBS combination but different 3HB phenotypes.

### Quantitative analysis of 3HB production of 15 selected strains under two light conditions

In order to properly analyze and compare the clone candidates from the preliminary screen, the 15 selected strains (**Figure 5**) were cultured in three independent replicates under two different growth conditions for a period of 14 days. The first condition (1% CO_2_ 50 μmol photons m^−2^s^−1^) represented the initial screening setup, while the second condition (3% CO_2_ 200 μmol photons m^−2^s^−1^) was selected for potentially enhancing 3HB productivity under higher light and carbon. All cultures were analyzed at three time points (days 8, 11 and 14) for cell growth and 3HB accumulation (**Figure 6; Supplementary Figures S5-S19**), and used for calculating the production levels and rates presented in **Table 3**. The growth of the cells was relatively uniform under each condition, and no obvious strain-specific differences or adverse effects were observed by the last sampling day 14. Under 3% CO_2_ and 200 μmol photons m^−2^s^−1^ light the exponential growth of the strains typically slowed down after the first week of cultivation (**Supplementary Figure S20**) after which the cultures maintained slow steady growth until d14 reaching OD_750nm_ 14-16. The first signs of bleaching were observed at d18, and by d20 all cultures had lost their green color. The 3HB accumulation for the strains varied from about 2 mgml^−1^ to almost 12 mgml^−1^ over the 14d sampling period, with six of the strains producing more than 5 mgml^−1^ 3HB in the culture medium (**Figure 6**; **Table 3**). Typically, the most efficient production was observed between the sampling points d11-d14, with the highest average daily production rates ranging between ∼1-3 gl^−1^d^−1^. Under the screening conditions, with 1% CO_2_ and 50 μmol photons m^−2^s^−1^ light, the cell growth was clearly slower and the 3HB production levels remained significantly lower, below 2 gl^−1^ for all the strains. As not all the possible RBS combinations were represented among the 15 analyzed strains, and the dataset was heavily biased by the non-random strain selection, it was not possible to reliably establish correlations between the RBS distribution and the recorded 3HB productivity (**Supplementary Figure S21**). Despite this limited information on RBS-context interactions, however, the data enabled us to demonstrate that different RBS combinations lead to varying levels of 3HB, and to identify promising strain candidates for further research.

**Figure 6:**
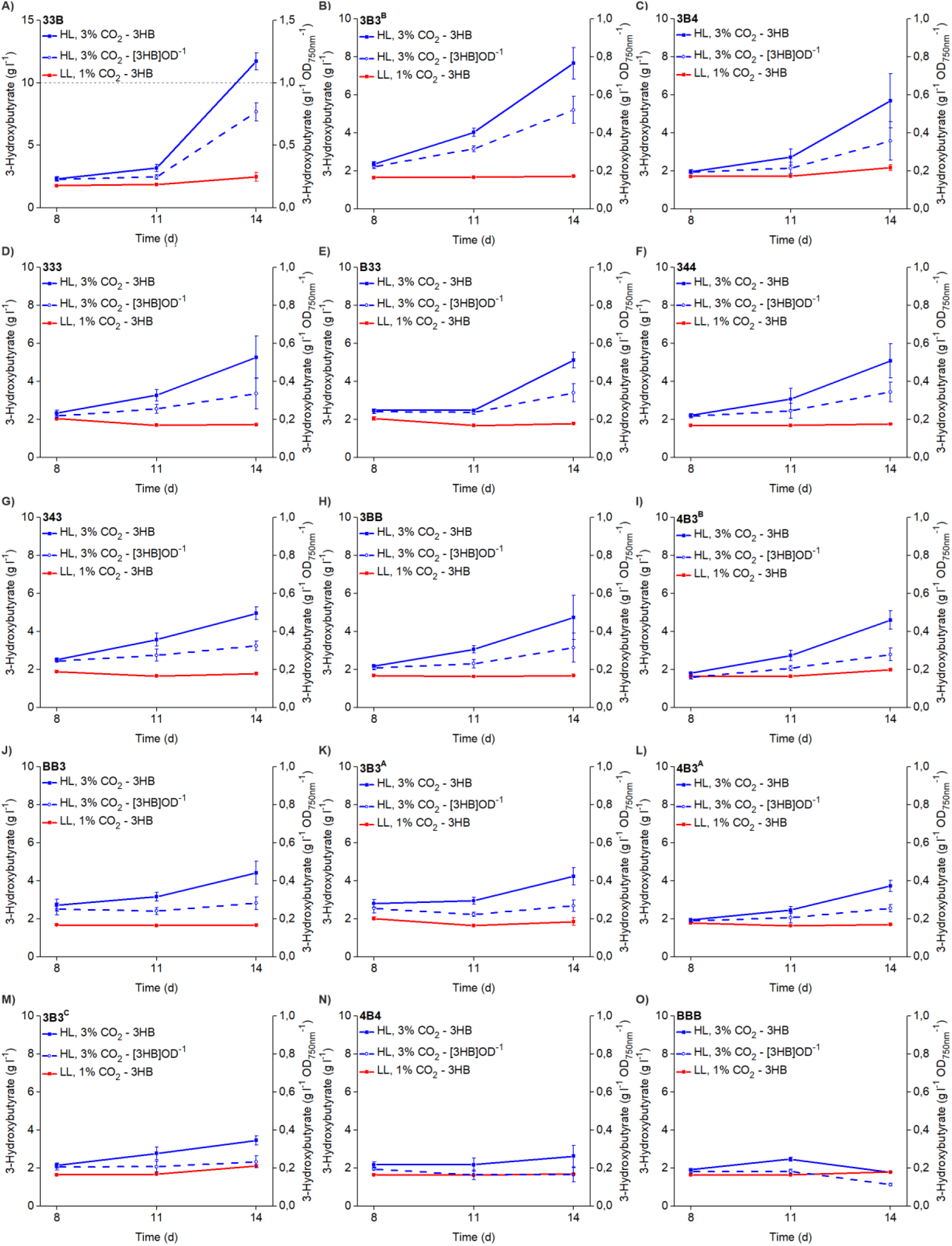
Comparison of the 15 selected *Synechocystis* strains based on 3HB production. Each strain was grown in three independent culture replicates for 14 days under two different conditions 3% CO_2_ 200 μmol photons m^−2^s^−1^ (HL) and 1% CO_2_ 50 μmol photons m^−2^s^−1^ (LL) in 20ml BG11 in 50ml Erlenmeyer flasks. The samples were collected on days 8, 11 and 14 and analyzed for 3HB accumulation. The solid lines represent the concentration of 3HB in the culture, while the dashed line is the value normalized to optical density OD_750nm_ ([3HB]OD^−1^). The averages and standard deviations have been calculated based on three independent culture replicates (n =3). To minimize variance from the GC-MS analysis, each sample was analyzed twice, and the values furthest from the total average were excluded. See complete data in **Supplementary Figures S5-S19**.

**Table 3:** Maximum 3HB accumulation and peak productivity (d11-d14) of the 15 selected *Synechocystis* strains in a two-week batch culture. The values have been calculated from the data presented in **Figure 6** from three independently conducted cultivations (n = 3) grown in 20 ml BG11 in 50 ml Erlenmeyer flasks under 3% CO_2_ 200 μmol photons m^−2^s^−1^. The seven strains selected for proteomics analysis are highlighted in red.

| Strain | Total 3HB<br>(g L<sup>-1</sup>) | 3HB productivity<br>d11-d14 (g L<sup>-1</sup>d<sup>-1</sup>) |
| --- | --- | --- |
| 33B | 11.72 | 2.85 |
| 3B3<sup>B</sup> | 7.66 | 1.21 |
| 3B4 | 5.69 | 0.99 |
| 333 | 5.27 | 0.67 |
| B33 | 5.13 | 0.68 |
| 344 | 5.09 | 0.67 |
| 343 | 4.96 | 0.46 |
| 3BB | 4.74 | 0.56 |
| 4B3<sup>B</sup> | 4.61 | 0.62 |
| BB3 | 4.43 | 0.42 |
| 3B3<sup>A</sup> | 4.24 | 0.43 |
| 4B3<sup>A</sup> | 3.72 | 0.42 |
| 3B3<sup>C</sup> | 3.46 | 0.23 |
| 4B4 | 2.63 | 0.15 |
| BBB | 1.78 | -0.23 |

### Quantitative proteomic analysis of the expressed 3HB pathway enzymes

Seven of the characterized strains, including the three highest 3HB producers, were selected for detailed comparison of the expression levels of TesB, PhaB and PhaA by targeted proteomics. The strains were cultivated in parallel in three independent replicates under 3% CO_2_ 200 μmol photons m^−2^s^−1^ and sampled at three time points on days 8, 11 and 14 as before. The quantitation of selected proteins was done by targeted LC-MS/MS profiling of three peptides per protein. The endogenous peptides were compared to internal heavy peptide standard reference, spiked in equal amounts to all samples, to ensure accurate representation of relative levels between PhaA, PhaB and TesB. The proteomic data was highly consistent between the replicates, and showed strain-specific differences in the total amounts and the ratios of PhaA, PhaB and TesB depending on different RBS combinations (**Figure 7; Supplementary Figures S22-S24**). Notably, within each strain the protein levels remained practically unchanged between the consecutive analytical time points (**Supplementary Figure S22**) indicating that each target protein maintained a strain-specific concentration equilibrium throughout the six-day sampling period. However, the protein levels did not have any profound effect on 3HB productivity at the first two time points, and the strain-specific differences in 3HB accumulation were mainly observed on day 14. (**Figure 6** vs **Supplementary Figure S22)**. This suggested that at the earlier phase of the batch culture the outcome was primarily determined by the metabolic flux to the substrate acetyl-CoA rather than the expression levels of TesB, PhaB and PhaA. Generally, the strains that produced 3HB more efficiently had higher overall content of all the three proteins as compared to the lower producers. The abundance of PhaA was consistently the highest of all the pathway proteins in all the strains, followed by TesB and PhaB in a decreasing order in the four best strains (**Figure 7 A-C**), while in the lower-producing strains the PhaB-TesB levels were more similar (**Figure 7 D-F; Supplementary Figures S22-S23**). Notably, in the top four strains, the differences between the individual target proteins appeared to be very small, and in many cases statistically insignificant (**Figure 7 A-C**; **Table 4; Supplementary Tables S2-S10**), indicating that even small variations in the relative ratios between the expression patterns may account for notable changes in the 3HB yields. As an example, the relative level of the first pathway enzyme PhaA appeared to systematically slightly lower in the best producer strain 33B in comparison the next best strains (**Supplementary Figures S22-S23**), yet we did not observe statistically significant differences in the calculated protein ratios **(Table 4; Supplementary Tables S11-S13)** even though the overall data quality and reproducibility were high.

**Figure 7:**
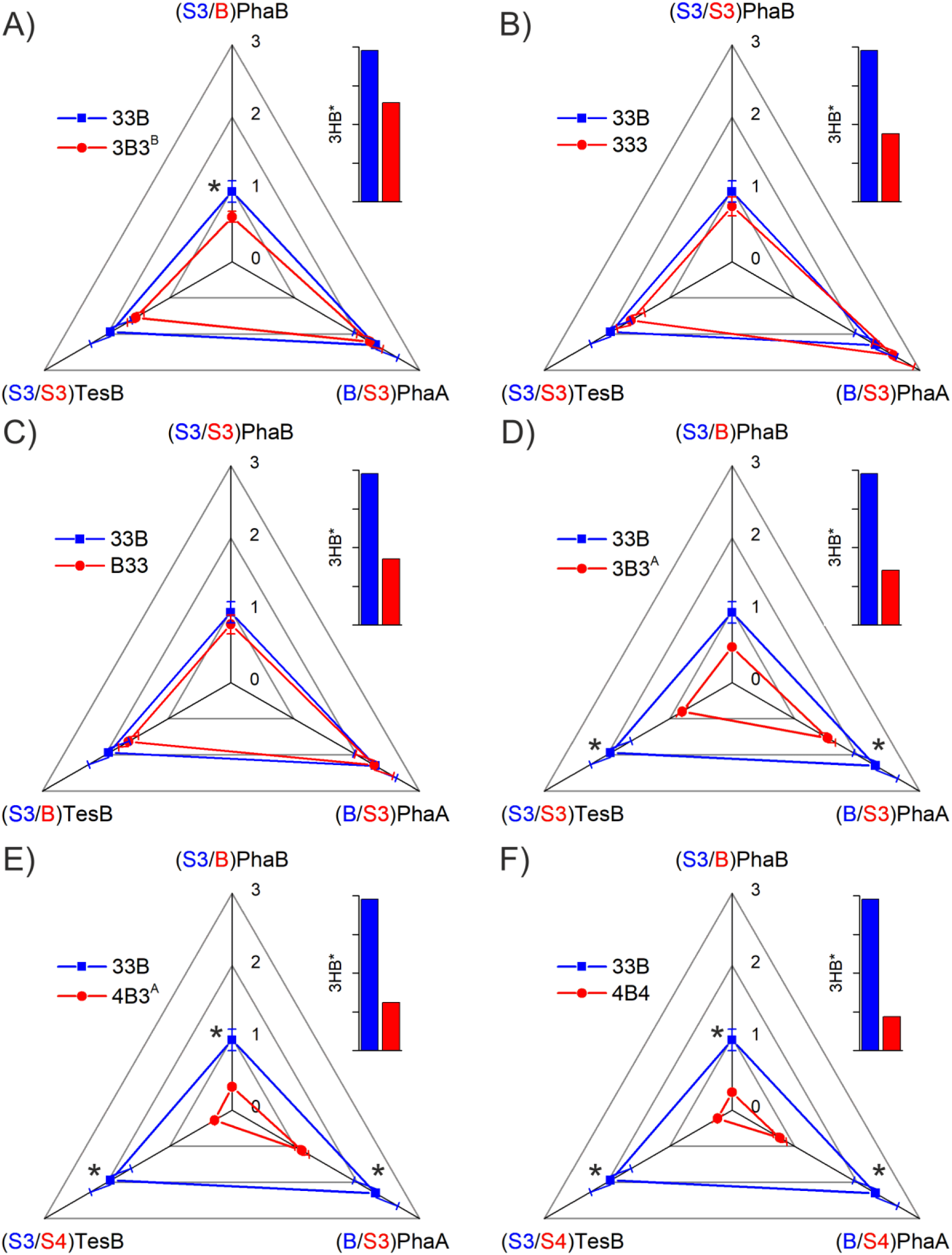
Radar plot illustrating the relative abundance of the 3HB pathway proteins TesB, PhaB and PhaA in the seven *Synechocystis* library strains selected for quantitative proteomic comparison. The protein profile of the highest-producing strain 33B (blue line) serves as a reference for comparing expression in **A)** 3B3^B^ **B)** 333 **C)** B33 **D)** 3B3^A^ **E)** 4B3^A^ **F)** 4B4. The corners of the radar plot correspond to the three overexpressed pathway proteins, with the RBSs indicated in text for each strain. The averages and standard deviations representing the abundances have been calculated based on all the proteomic data from the three sampling points d8, d11, d14 combined (n > 9). Statistically significant (two-tailed Student’s t-tests p < 0.05) differences in protein abundance between the two strains are marked with * for each protein in question. The bar chart next to each radar plot illustrates the relative differences in the 3HB accumulation between the strains measured at d14.

**Table 4:** Calculated statistical differences in the abundance of TesB, PhaA and PhaB between the highest 3HB producer 3BB in comparison to the six other strains subjected to proteomic analysis. A positive standardized difference d indicates that the strain 33B has a larger value or ratio, whereas a negative d indicates that the comparator strain has larger value or ratio. Blue: significant statistical difference (p < 0.05) in the protein abundance between 33B and the comparator strain. Red: no statistical significance (p > 0.05) in the protein abundance between 33B and the comparator strain. Purple: no statistical significance, but medium effect in the standardized difference between the strain 33B and the comparator strain (∣d∣ > 0.5). The values have been calculated based on all the proteomic data from the three sampling points d8, d11, d14 combined (n > 9). See **Supplementary Tables S2-S13** for details.

| Comparator Strain | TesB<br>p / (d) | PhaA<br>p / (d) | PhaB<br>p / (d) | TesB:PhaA<br>p / (d) | TesB:PhaB<br>p / (d) | PhaA:PhaB<br>p / (d) |
| --- | --- | --- | --- | --- | --- | --- |
| **3B3<sup>B</sup>** | 0.246<br>(0.274) | 0.708<br>(0.040) | 0.011<br>(0.884) | 0.420<br>(0.568) | 0.420<br>(-0.583) | 0.080<br>(-0.459) |
| **333** | 0.022<br>(0.205) | 0.897<br>(-0.126) | 0.522<br>(0.430) | 0.518<br>(0.521) | 0.518<br>(-0.556) | 0.193<br>(-0.332) |
| **B33** | 0.602<br>(0.216) | 0.689<br>(0.007) | 0.651<br>(0.364) | 0.699<br>(0.450) | 0.699<br>(-0.498) | 0.415<br>(-0.204) |
| **3B3<sup>A</sup>** | 0.000<br>(0.926) | 0.136<br>(0.367) | 0.000<br>(1.273) | 0.223<br>(0.657) | 0.230<br>(-0.285) | 0.346<br>(-0.219) |
| **4B3<sup>A</sup>** | 0.000<br>(1.425) | 0.512<br>(0.619) | 0.000<br>(1.828) | 0.000<br>(1.401) | 0.000<br>(0.240) | 0.065<br>(-0.456) |
| **4B4** | 0.000<br>(1.468) | 0.020<br>(0.835) | 0.000<br>(2.107) | 0.001<br>(1.218) | 0.001<br>(0.246) | 0.234<br>(-0.287) |

## Discussion

This work focused on the photoautotrophic production of the soluble bioplastic precursor 3HB in engineered cyanobacterium *Synechocystis*, and demonstrated a profound increase in the efficiency through enzyme-level pathway optimization. We specifically targeted a secreted product that can be harvested from the culture without breaking the cells, with a potential for developing continuous bioproduction systems with higher production rates and lower unit costs as compared to batch cultures [46]. Secretion also prevents end-product inhibition and shifts the pathway equilibrium from the reaction precursor (acetyl-CoA) towards the product [47], which is a biosynthetic advantage for producing 3HB as compared to the polymeric form PHB. The 3HB pathway offered a good platform for translational tuning because the three engineered steps have a clear combined effect on the flux [29] [30], while the productivity is not critically limited by substrate availability at the production phase. Our results show that the expression of the three 3HB pathway enzymes PhaA, PhaB and TesB **(Figure 1)** at different levels under the regulation of three alternative RBSs allowed us to build a strain library (**Figure 3**) with a broad range of 3HB production efficiencies (**Figure 5; Supplementary Figures S5-S19**). This led to the identification of several strains that accumulated 3HB efficiently at gram per liter levels (**Table 3**) that are higher than previously reported in any photoautotrophic host. Multi-gene translational pathway optimization has not been extensively explored in cyanobacterial engineering, and this work provides a successful example of the use and potential of the strategy, which could possibly be applied as a general tool for photoautotrophic strain development.

Earlier strategies for increasing 3HB productivity in cyanobacteria have based on the identification and stepwise reinforcement of the limiting steps in the heterologous pathway [29] [30]. Wang et al demonstrated that acetoacetyl-CoA reductase PhaB was a key bottleneck in their engineered strain, and that the optimization of the RBS lead to higher enzyme activity along with increased 3HB productivity to 1.84 gl^−1^ within 10 days in *Synechocystis* [29]. Ku and Lan enhanced their pathway by introducing an additional ATP-driven acetoacetyl-CoA synthase NphT7 to catalyze the conversion of acetyl-CoA in parallel to PhaA in *Synechococcus elongatus* PCC 7942 [30]. In combination with the expression of additional thioesterases to accelerate the release of CoA in the last step, this resulted in the cumulative 3HB titer 1.2 gl^−1^ over a 28-day culture period. These studies clearly illustrate that the outcome is dictated by the balance between the successive pathway steps, and that flux limitation changes every time when a specific step is optimized. Here, we took a different approach; rather than adjusting the biosynthetic steps one by one, we altered the expression patterns of the three pathway enzymes simultaneously by using three alternative RBSs for each gene. As the outcome, six of the engineered strains characterized here (**Figure 6**) accumulated over 5 gl^−1^ 3HB in the culture medium over a two-week batch cultivation, with the highest producing strain exceeding total 11 gl^−1^ (**Table 3**). While direct quantitative comparison between the different experimental setups is challenging, this represents a clear improvement in the reported photoautotrophic 3HB production levels, and is comparable to the best engineered heterotrophic systems reaching up to 16 gl^−1^ in *E. coli* [48] and 12 gl^−1^ in *S. cerevisiae* [24] when using carbohydrates as substrates. In line with earlier 3HB accumulation profiles, the highest productivity was systematically observed at the end of the two-week batch cultivation period, following the exponential growth phase. This coincides with the depletion of nitrogen and phosphate from the medium which is known to effectively induce 3HB production [28] by redirecting the cellular carbon flux from biomass formation towards acetyl-CoA and the 3HB pathway when growth is limited [29]. For the three most efficient strains the maximum production rates at this phase were about 1 gl^−1^ d^−1^, 1.2 gl^−1^ d^−1^ and 2.9 gl^−1^ d^−1^ 3HB, as calculated for the batch culture days 11-14, as compared to previously reported peak productivities ∼0,3 gl^−1^ d^−1^ [29] and ∼0.1 gl^−1^ d^−1^ [30].

In addition to the improved flux at the 3HB pathway level, the strain performance was also affected by other factors including the use of replicative plasmid with a higher copy number compared to chromosomal integration [36], implementation of efficient RBSs that derive from highly expressed genes in *Synechocystis* [32] and high-carbon and high light conditions that favor production in extended batch cultivation. In comparison to PHB, although the highest accumulation levels in cyanobacteria have reached up to 81% [10] or even 85% [49] of the dry cell weight, the productivity of PHB has remained only at milligram levels per liter per day [6]. This suggests that some of the key constraints associated with intracellular polymer formation could be avoided by producing derivatives that are secreted out from the cell. Notably, life cycle assessment on photoautotrophic PHB production has suggested that the light-driven systems utilizing CO_2_ as the carbon source would need effective recovery at levels above 0.8 gl^−1^ d^−1^ [50] to reach commercial market price levels. Even though product harvest and upscale have not yet been addressed, the achieved 3HB levels are in the desired order of magnitude. While this is a promising start towards developing competitive continuous biotechnological systems for bioplastic precursors, there are still open fundamental questions regarding factors that affect the target protein expression and pathway flux under varying metabolic states and cultivation conditions.

The efficiency of protein expression is determined at several regulatory levels in vivo, to which the RBSs generally only have a partial relative effect. The efficiency of transcription determines the amount of available mRNA template for translation, while posttranscriptional modifications in cyanobacteria can critically affect transcript stability and ribosome accessibility [51] [52]. These native processes have been studied in several cyanobacterial contexts, and it appears that in many cases the system is directly or indirectly regulated at the transcript level, often showing a correlation between mRNA and protein abundance [53] [54]. For example, extensive analysis of wild-type *Synechocystis* under long-term day-night cycling has shown that there is a low degree of translational regulation determining the native protein quantities *in vivo*, and that the oscillations rather follow changes at the transcript level [54]. However, as demonstrated here, in engineered systems the expression can also be effectively regulated at the level of translation by using alternative RBSs, with a consequent direct catalytic impact on the 3HB production efficiency (**Figure 6**, **Table 3**). Controlled by a single promoter, the three target proteins PhaA, PhaB and TesB were shown to have strain-specific expression patterns, with relatively constant intracellular levels from one time point to another in the non-steady state batch culture (**Supplementary Figures S22-S24**). The comparison revealed that quite substantial differences in 3HB productivity (**Figure 5**; **Table 3**) could sometimes be attributed to very subtle variation in the pathway protein levels (**Figure 7; Supplementary Figure S23**). This suggests that small consistent differences in enzyme abundance (that may be difficult to quantitate even with accurate proteomic methods) can lead to substantial cumulative catalytic effects over time that are reflected on the production phenotype. For example, although the calculated differences appear insignificant (**Table 4**), the expression data on the highest-producing strains supports the view that increased relative amount of PhaA over PhaB and TesB may be linked to lowered 3HB yield (**Figure 7 A-C**) as discussed before [28] [30]. Besides the combined effect of the protein quantity and the enzyme-specific catalytic kinetic properties, the outcome is also clearly affected by the culture parameters (**Figure 5** vs. **Figure 6**). In the 3HB strains these determine, for example, the metabolic flux to the acetyl-CoA precursor and the availability of PhaB co-substrate NADPH under different experimental conditions. For this reason, the most optimal RBS combination for 3HB production may vary from one cultivation setup to another. As the current study was conducted in batch mode, further research is necessary to assess the strain performance under carefully adjusted continuous steady-state and fed-batch systems, where the metabolic equilibria determining the availability of acetyl-CoA and the effect of enzyme levels may be different.

It is clear that the target protein expression patterns can be profoundly altered by using alternative RBSs in vivo. However, it is difficult to quantitate the relative RBS efficiencies or univocally assign the best RBS for a particular target gene because of the significant sequence context-dependence influencing expression [32] [35] [33] (**Supplementary Figure S23-S24**). Although the molecular factors behind varying translation patterns are not fully understood [see also [34]], they are expected to involve mRNA 5’ UTR secondary structures and possible association with specific mRNA binding-proteins (RBPs) [55] that can affect transcript stability, ribosome accessibility or start codon recognition. In addition, high ribosome occupancy on multicistronic transcripts, as here for PhaA, PhaB and TesB, may also cause indirect steric effects on the translation of adjacent proteins [56]. This means that the translation efficiency of a given RBS-gene pair is not necessarily uniform but influenced by the translation levels of the other two pathway proteins in the operon, which may explain some of the observed variance between the 3HB constructs (**Supplementary Figure S23-S24**). The systematic comparison of the RBS effects in the current work was further constrained by the lack of expression data on all the 27 combinations, and changes associated with spontaneous mutations [see also [57] [58] [59] [60]] that are the likely cause for some of the observed phenotypes. Even though understanding these interactions would be important for rational engineering, the work here demonstrates how the combinatorial approach can be used to simultaneously tune the intracellular levels of multiple enzymes to improve productivity, without knowing all the overlapping factors that influence the outcome. Marked effects were achieved with only three different RBSs in a three-gene operon, but the process could be further enhanced by using a larger set of the control elements [32] for tuning the intracellular target protein levels for optimal production.

Despite the development of strain-specific synthetic biology tools and access to fast routine gene synthesis, construct assembly is still one of the major time-consuming bottlenecks in cyanobacterial engineering. Applications that depend on the generation of numerous construct variants, as in the case of the 3HB pathways in this work, would benefit from more efficient preparative workflows that improve throughput. While the modular cloning system implemented here allowed the assembly of the three-gene 3HB constructs with different RBSs for adjusting expression levels, the approach is relatively laborious and requires multiple iterative rounds of subcloning. As an alternative, various multi-part DNA assembly strategies that enable the generation increasingly complex genetic constructs with less effort have been adapted for cyanobacterial engineering and translational tuning [35] [61] [62]. The most recent Golden Gate –based system that has been specifically designed for this purpose [63] enables the assembly of transformation-ready multicistronic expression constructs in a single one-pot reaction. This platform is based on the same RBS library as used in the current work, and well applicable for the combinatorial modular assembly of multiple three-gene expression construct variants in parallel. Together with case-studies that demonstrate the advantage of translation level optimization in different contexts, such high-throughput assembly tools lower the threshold for using RBS-based optimization for protein overexpressing in cyanobacteria, and improve prospects in future bioprocess development.

## Supporting information

Supplementary information for Kakko et al

## Summary

This work demonstrates the use of translational tuning to balance the expression levels of three consecutive target pathway enzymes in engineered cyanobacterial host *Synechocystis*, enabling significantly enhanced photoautotrophic production of secreted bioplastic precursor 3HB from CO_2_ at a gram per liter range. The results have important general implications for metabolic engineering, and highlight the potential of adjusting the amounts of target proteins in the cell without prior knowledge of relative expression efficiencies, enzyme kinetic properties or protein-specific degradation rates.

## Acknowledgements & funding

Jane and Aatos Erkko Foundation. Jenny and Antti Wihuri Foundation. The Finnish Foundation for Technology Promotion. Kone Foundation.

