## Supplementary information for Kakko et al for "Optimizing photoautotrophic production of soluble 3-hydroxybutyrate in *Synechocystis* sp. PCC 6803 through combinatorial translational tuning"

**Supplementary Figure S1**: Customized LED system for cyanobacterial cultures……………….….….……p.2
**Supplementary Table S1**: Proteomics sample information on TesB, PhaA and PhaB…………………….... p.3
**Supplementary Figure S2**: Nucleotide sequences of the ordered *tesB, phaA* and *phaB* constructs………..p.4-5
**Supplementary Figure S10**: Strain 344 complete datasets on 3HB production and growth……………...…p.13
**Supplementary Figure S11**: Strain 343 complete datasets on 3HB production and growth……………...…p.14
**Supplementary Figure S12**: Strain 3BB complete datasets on 3HB production and growth……………….…p.15
**Supplementary Figure S13**: Strain 4B3[B] complete datasets on 3HB production and growth……………….…p.16
**Supplementary Figure S14**: Strain BB3 complete datasets on 3HB production and growth……………...…p.17
**Supplementary Figure S18**: Strain 4B4 complete datasets on 3HB production and growth……………...…p.21
**Supplementary Figure S19**: Strain BBB complete datasets on 3HB production and growth……………...…p.22
**Supplementary Figure S20**: Representative growth profile measured every second day for 20 d………..…p.23
**Supplementary Figure S21.** Box and whiskers plot; gene-RBS combinations vs. 3HB…………………….…p.24
**Supplementary Figure S22**. Relative abundance of TesB, PhaB and PhaA at three time time points………p.25
**Supplementary Figure S23**. The averaged total abundance of TesB, PhaB and PhaA…………………....…p.26
**Supplementary Figure S24.** Box and whiskers plot of TesB, PhaA and PhaB abundance……………..……p.27
**Supplementary Table S2**. Mann-Whitney U test comparing TesB abundance between strains…………..…p.28
**Supplementary Table S3**. Mann-Whitney U test comparing PhaA abundance between strains…………..…p.29
**Supplementary Table S4**. Mann-Whitney U test comparing PhaB abundance between strains…………..…p.30
**Supplementary Table S5**. Approximate z-test for log-scale TesB:PhaA ratio between strains…………..…p.31
**Supplementary Table S6**. Approximate z-test for log-scale TesB:PhaB ratio between strains…………..…p.32
**Supplementary Table S7**. Approximate z-test for log-scale PhaA:PhaB ratio between strains……….….…p.33
**Supplementary Table S8**. Cohen's d effect sizes for TesB abundance between strains.…………………..…p.34
**Supplementary Table S9**. Cohen's d effect sizes for PhaA abundance between strains……………….…..…p.35
**Supplementary Table S10**. Cohen's d effect sizes for PhaB abundance between strains.……………….…..…p.36
**Supplementary Table S11**. Cohen's d effect sizes for log-scale TesB:PhaA ratio between strains……..……p.37
**Supplementary Table S12**. Cohen's d effect sizes for log-scale TesB:PhaB ratio between strains. ……...…p.38
**Supplementary Table S13**. Cohen's d effect sizes for log-scale PhaA:PhaB ratio between strains……..……p.39

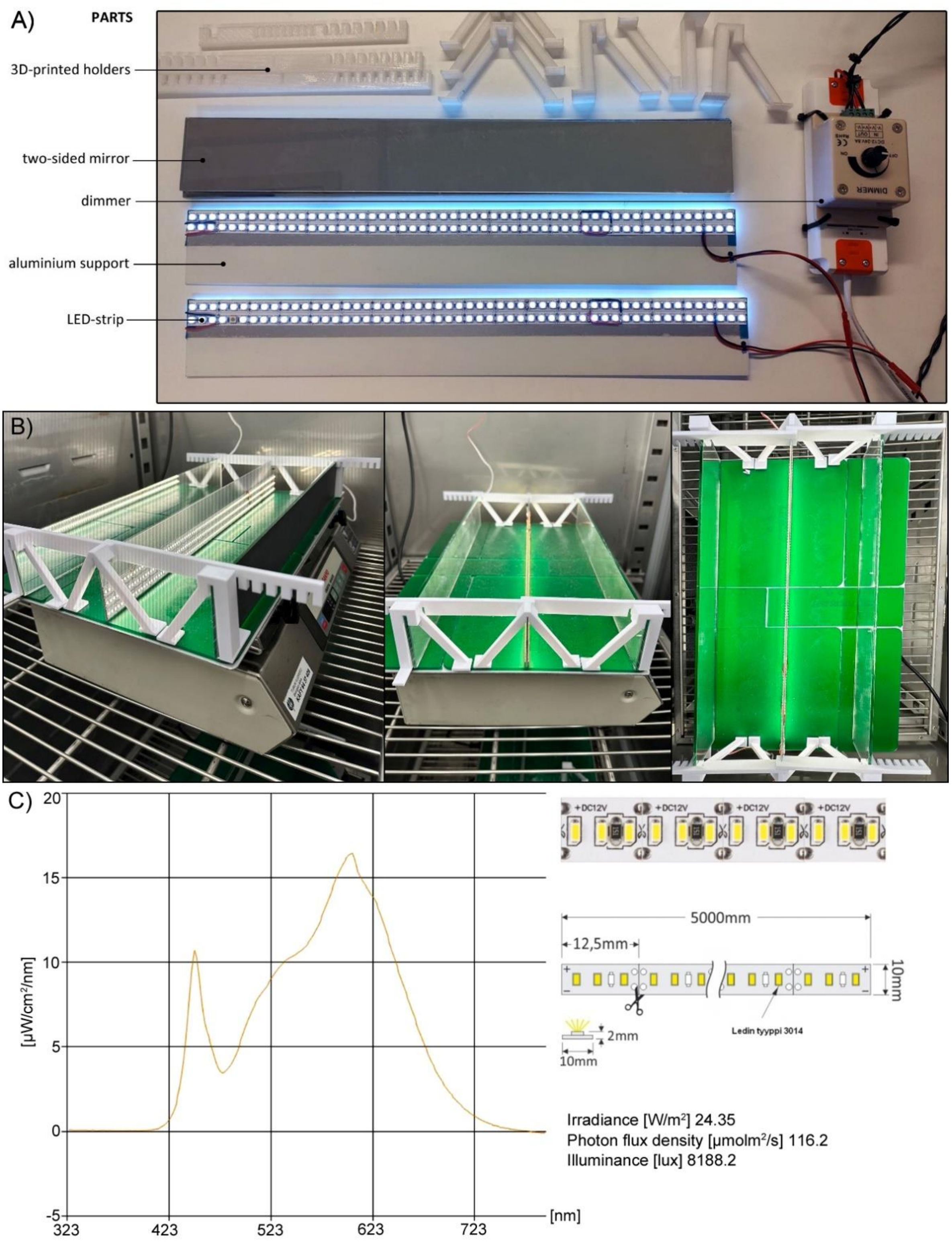

**Supplementary Figure S1**: Custom-built LED setup to ensure uniform lightning for the parallel cyanobacterial cultures. **A)** The system was compiled from LED strips and their control driver, mirrors, aluminum strips and 3D-printed stands **B)** The lights were fixed in place on a Sanyo orbital shaker with sticky pads, and placed inside a POL-EKO Apartura culture cabinet to control temperature and $CO_2$ concentration. The Erlenmeyer culture flasks were placed at equal distances between the LEDs on one side and a mirror on the other side. The light intensity in was measured from the middle inside the flasks, and set to the desired value using the dimmer. **C)** The LEDs were commercial fully dimmable high-density diode strips (12 V DC, 90 W total power) emitting warm-white light (3000 K; CRI $\geq$ 80) with luminous flux of 12,000 lm. The LEDs were rated IP20 for indoor dry-area use, compliant with RoHS standards.

**Supplementary Table S1**: List of peptide sequences (light-endogenous and heavy-spiked reference) and corresponding parent ions (m/z) used in scheduled PRM mass spectrometry method for quantification of three engineered proteins TesB (P0AGG2), PhaB (P14697) and PhaA (P14611). The data for parent ions were registered in positive ionization mode with normalized collision energy (NCE) of 27V in time windows of two minutes. Modified amino acids in heavy references are marked with bold font.

| Mass [m/z] | CS [z] | Polarity | Start [min] | End [min] | NCE | Peptide sequence | Protein (Uniprot ID) |
|---|---|---|---|---|---|---|---|
| 777.9299 | 2 | Positive | 45.6 | 47.6 | 27 | FDEEIVPVLIPQR (light) | P14611 |
| 782.9341 | 2 | Positive | 45.6 | 47.6 | 27 | FDEEIVPVLIPQR (heavy) | ref |
| 578.3528 | 2 | Positive | 42.41 | 44.41 | 27 | ELGLTPLATIK (light) | P14611 |
| 582.3599 | 2 | Positive | 42.41 | 44.41 | 27 | ELGLTPLATIK (heavy) | ref |
| 511.2511 | 2 | Positive | 16.28 | 18.28 | 27 | SYANAGVDPK (light) | P14611 |
| 515.2582 | 2 | Positive | 16.28 | 18.28 | 27 | SYANAGVDPK (heavy) | ref |
| 537.2777 | 2 | Positive | 27.19 | 29.19 | 27 | VMGMGPVPASK (light) | P14611 |
| 541.2848 | 2 | Positive | 27.19 | 29.19 | 27 | VMGMGPVPASK (heavy) | ref |
| 557.2771 | 2 | Positive | 13.42 | 15.42 | 27 | VVAGC[+57.021464]GPNSPR (light) | P14697 |
| 562.2812 | 2 | Positive | 13.42 | 15.42 | 27 | VVAGC[+57.021464]GPNSPR (heavy) | ref |
| 579.8275 | 2 | Positive | 24.64 | 26.64 | 27 | IVNISSVNGQK (light) | P14697 |
| 583.8346 | 2 | Positive | 24.64 | 26.64 | 27 | IVNISSVNGQK (heavy) | ref |
| 420.7813 | 2 | Positive | 27.86 | 29.86 | 27 | IVATIPVK (light) | P14697 |
| 424.7884 | 2 | Positive | 27.86 | 29.86 | 27 | IVATIPVK (heavy) | ref |
| 673.8876 | 2 | Positive | 37.42 | 39.42 | 27 | KPIIYDVETLR (light) | P0AGG2 |
| 678.8917 | 2 | Positive | 37.42 | 39.42 | 27 | KPIIYDVETLR (heavy) | ref |
| 427.1936 | 2 | Positive | 13.5 | 15.5 | 27 | DGNSFSAR (light) | P0AGG2 |
| 432.1977 | 2 | Positive | 13.5 | 15.5 | 27 | DGNSFSAR (heavy) | ref |
| 522.2594 | 2 | Positive | 21.58 | 23.58 | 27 | ANGSVPDDLR (light) | P0AGG2 |
| 527.2636 | 2 | Positive | 21.58 | 23.58 | 27 | ANGSVPDDLR (heavy) | ref |

### A) *tesB*-CmR

ATGCATAGTCAGGCGCTAAAAAATTTACTGACATTGTTAAATCTGGAAAAAATTGAGGAAGGACTCTTTCGCGGCCAGAGTGA
AGATTTAGGTTTACGCCAGGTGTTTGGCGGCCAGGTCGTGGGTCAGGCCTTGTATGCTGCAAAAGAGACCGTCCCTGAAGAGC
GGCTGGTACATTCGTTTCACAGCTACTTTCTTCGCCCTGGCGATAGTAAGAAGCCGATTATTTATGATGTCGAAACGCTGCGT
GACGGTAACAGCTTCAGCGCCCGCCGGGTTGCTGCTATTCAAAACGGCAAACCGATTTTTTATATGACTGCCTCTTTCCAGGC
ACCAGAAGCGGGTTTCGAACATCAAAAAACAATGCCGTCCGCGCCAGCGCCTGATGGCCTCCCTTCGGAAACGCAAATCGCCC
AATCGCTGGCGCACCTGCTGCCGCCAGTGCTGAAAGATAAATTCATCTGCGATCGTCCGCTGGAAGTCCGTCCGGTGGAGTTT
CATAACCCACTGAAAGGTCACGTCGCAGAACCACATCGTCAGGTGTGGATCCGCGCAAATGGTAGCGTGCCGGATGACCTGCG
CGTTCATCAGTATCTGCTCGGTTACGCTTCTGATCTTAACTTCCTGCCGGTAGCTCTACAGCCGCACGGCATCGGTTTTCTCG
AACCGGGGATTCAGATTGCCACCATTGACCATTCCATGTGGTTCCATCGCCCGTTTAATTTGAATGAATGGCTGCTGTATAGC
GTGGAGAGCACCTCGGCGTCCAGCGCACGTGGCTTTGTGCGCGGTGAGTTTTATACCCAAGACGGCGTACTGGTTGCCTCGAC
CGTTCAGGAAGGGGTGATGCGTAATCACAATTAAGCTAGCGTTGATCGGGCACGTAAGAGGTTCCAACTTTCACCATAATGAA
ATAAGATCACTACCGGGCGTATTTTTTGAGTTATCGAGATTTTCAGGAGCTAAGGAAGCTAAAATGGAGAAAAAAATCACGGG
ATATACCACCGTTGATATATCCCAATGGCATCGTAAAGAACATTTTGAGGCATTTCAGTCAGTTGCTCAATGTACCTATAACC
AGACCGTTCAGCTGGATATTACGGCCTTTTTAAAGACCGTAAAGAAAAATAAGCACAAGTTTTATCCGGCCTTTATTCACATT
CTTGCCCGCCTGATGAACGCTCACCCGGAGTTTCGTATGGCCATGAAAGACGGTGAGCTGGTGATCTGGGATAGTGTTCACCC
TTGTTACACCGTTTTCCATGAGCAAACTGAAACGTTTTCGTCCCTCTGGAGTGAATACCACGACGATTTCCGGCAGTTTCTCC
ACATATATTCGCAAGATGTGGCGTGTTACGGTGAAAACCTGGCCTATTTCCCTAAAGGGTTTATTGAGAATATGTTTTTGTC
TCAGCCAATCCCTGGGTGAGTTTCACCAGTTTTGATTTAAACGTGGCCAATATGGACAACTTCTTCGCCCCCGTTTTCACGAT
GGGCAAATATTATACGCAAGGCGACAAGGTGCTGATGCCGCTGGCGATCCAGGTTCATCATGCCGTTTGTGATGGCTTCCATG
TCGGCCGCATGCTTAATGAATTACAACAGTACTGTGATGAGTGGCAGGGCGGGGCGTAATAAGCTAGCGCGGCCGCTCGAG

### B) phaB-CmR

ATGCATACTCAGCGCATTGCGTATGTGACCGGCGGCATGGGTGGTATCGGAACCGCCATTTGCCAGCGGCTGGCCAAGGATGG
CTTTCGTGTGGTGGCCGGTTGCGGCCCCAACTCGCCGCGCCGCGAAAAGTGGCTGGAGCAGCAGAAGGCCCTGGGCTTCGATT
TCATTGCCTCGGAAGGCAATGTGGCTGACTGGGACTCGACCAAGACCGCATTCGACAAGGTCAAGTCCGAGGTCGGCGAGGTT
GATGTGCTGATCAACAACGCCGGTATCACCCGCGACGTGGTGTTCCGCAAGATGACCCGCGCCGACTGGGATGCGGTGATCGA
CACCAACCTGACCTCGCTGTTCAACGTCACCAAGCAGGTGATCGACGGCATGGCCGACCGTGGCTGGGGCCGCATCGTCAACA
TCTCGTCGGTGAACGGGCAGAAGGGCCAGTTCGGCCAGACCAACTACTCCACCGCCAAGGCCGGCCTGCATGGCTTCACCATG
GCACTGGCGCAGGAAGTGGCGACCAAGGGCGTGACCGTCAACACGGTCTCTCCGGGCTATATCGCCACCGACATGGTCAAGGC
GATCCGCCAGGACGTGCTCGACAAGATCGTCGCGACGATCCCGGTCAAGCGCCTGGGCCTGCCGGAAGAGATCGCCTCGATCT
GCGCCTGGTTGTCGTCGGAGGAGTCCGGTTTCTCGACCGGCGCCGACTTCTCGCTCAACGGCGGCCTGCATATGGGCTGAGCT
AGCGTTGATCGGGCACGTAAGAGGTTCCAACTTTCACCATAATGAAATAAGATCACTACCGGGCGTATTTTTTGAGTTATCGA
GATTTTCAGGAGCTAAGGAAGCTAAAATGGAGAAAAAAATCACGGGATATACCACCGTTGATATATCCCAATGGCATCGTAAA
GAACATTTTGAGGCATTTCAGTCAGTTGCTCAATGTACCTATAACCAGACCGTTCAGCTGGATATTACGGCCTTTTTAAAGAC
CGTAAAGAAAAATAAGCACAAGTTTTATCCGGCCTTTATTCACATTCTTGCCCGCCTGATGAACGCTCACCCGGAGTTTCGTA
TGGCCATGAAAGACGGTGAGCTGGTGATCTGGGATAGTGTTCACCCTTGTTACACCGTTTTCCATGAGCAAACTGAAACGTTT
TCGTCCCTCTGGAGTGAATACCACGACGATTTCCGGCAGTTTCTCCACATATATTCGCAAGATGTGGCGTGTTACGGTGAAAA
CCTGGCCTATTTCCCTAAAGGGTTTATTGAGAATATGTTTTTGTCTCAGCCAATCCCTGGGTGAGTTTCACCAGTTTTGATT
TAAACGTGGCCAATATGGACAACTTCTTCGCCCCCGTTTTCACGATGGGCAAATATTATACGCAAGGCGACAAGGTGCTGATG
CCGCTGGCGATCCAGGTTCATCATGCCGTTTGTGATGGCTTCCATGTCGGCCGCATGCTTAATGAATTACAACAGTACTGTGA
TGAGTGGCAGGGCGGGGCGTAATAAGCTAGCGCGGCCGCTCGAG

### C) *phaA*-CmR

ATGCATACTGACGTTGTCATCGTATCCGCCGCCCGCACCGCGGTCGGCAAGTTTGGCGGCTCGCTGGCCAAGATCCCGGCACC
GGAACTGGGTGCCGTGGTCATCAAGGCCGCGCTGGAGCGCGCCGGCGTCAAGCCGGAGCAGGTGAGCGAAGTCATCATGGGCC
AGGTGCTGACCGCCGGTTCGGGCCAGAACCCCGCACGCCAGGCCGCGATCAAGGCCGGCCTGCCGGCGATGGTGCCGGCCATG
ACCATCAACAAGGTGTGCGGCTCGGGCCTGAAGGCCGTGATGCTGGCCGCCAACGCGATCATGGCGGGCGACGCCGAGATCGT
GGTGGCCGGCGGCCAGGAAAACATGAGCGCCGCCCCGCACGTGCTGCCGGGCTCGCGCGATGGTTTCCGCATGGGCGATGCCA
AGCTGGTGGACACCATGATCGTGGACGGCCTGTGGGACGTGTACAACCAGTACCACATGGGCATCACCGCCGAGAACGTGGCC
AAGGAATACGGCATCACACGCGAGGCGCAGGATGAGTTCGCCGTCGGCTCGCAGAACAAGGCCGAAGCCGCGCAGAAGGCCGG
CAAGTTTGACGAAGAGATCGTCCCGGTGCTGATCCCGCAGCGCAAGGGCGACCCGGTGGCCTTCAAGACCGACGAGTTCGTGC
GCCAGGGCGCCACGCTGGACAGCATGTCCGGCCTCAAGCCCGCCTTCGACAAGGCCGGCACGGTGACCGCGGCCAACGCCTCG
GGCCTGAACGACGGCGCCGCCGCGGTGGTGGTGATGTCGGCGGCCAAGGCCAAGGAACTGGGCCTGACCCCGCTGGCCACGAT
CAAGAGCTATGCCAACGCCGGTGTCGATCCCAAGGTGATGGGCATGGGCCCGGTGCCGGCCTCCAAGCGCGCCCTGTCGCGCG
CCGAGTGGACCCCGCAAGACCTGGACCTGATGGAGATCAACGAGGCCTTTGCCGCGCAGGCGCTGGCGGTGCACCAGCAGATG
GGCTGGGACACCTCCAAGGTCAATGTGAACGGCGGCGCCATCGCCATCGGCCACCCGATCGGCGCGTCGGGCTGCCGTATCCT
GGTGACGCTGCTGCACGAGATGAAGCGCCGTGACGCGAAGAAGGGCCTGGCCTCGCTGTGCATCGGCGGCGGCATGGGCGTGG

CGCTGGCAGTCGAGCGCAAATAAGCTAGCGTTGATCGGGCACGTAAGAGGTTCCAACTTTCACCATAATGAAATAAGATCACT
ACCGGGCGTATTTTTTGAGTTATCGAGATTTTCAGGAGCTAAGGAAGCTAAAATGGAGAAAAAAATCACGGGATATACCACCG
TTGATATATCCCAATGGCATCGTAAAGAACATTTTGAGGCATTTCAGTCAGTTGCTCAATGTACCTATAACCAGACCGTTCAG
CTGGATATTACGGCCTTTTTAAAGACCGTAAAGAAAAATAAGCACAAGTTTTATCCGGCCTTTATTCACATTCTTGCCCGCCT
GATGAACGCTCACCCGGAGTTTCGTATGGCCATGAAAGACGGTGAGCTGGTGATCTGGGATAGTGTTCACCCTTGTTACACCG
TTTTCCATGAGCAAACTGAAACGTTTTCGTCCCTCTGGAGTGAATACCACGACGATTTCCGGCAGTTTCTCCACATATATTCG
CAAGATGTGGCGTGTTACGGTGAAAACCTGGCCTATTTCCCTAAAGGGTTTATTGAGAATATGTTTTTTGTCTCAGCCAATCC
CTGGGTGAGTTTCACCAGTTTTGATTTAAACGTGGCCAATATGGACAACTTCTTCGCCCCCGTTTTCACGATGGGCAAATATT
ATACGCAAGGCGACAAGGTGCTGATGCCGCTGGCGATCCAGGTTCATCATGCCGTTTGTGATGGCTTCCATGTCGGCCGCATG
CTTAATGAATTACAACAGTACTGTGATGAGTGGCAGGGCGGGGCGTAATAAGCTAGCGCGGCCGCTCGAG

**Supplementary Figure S2**: **The DNA sequences of the synthetic genes A)** *tesB*, **B)** *phaB* and **C)** *phaB* ordered as CmR fusions for the pathway assembly. The target gene sequences are shown in black font, the spacer sequences in grey font and the CmR in grey highlight. The restriction sites are indicated in coloured highlight: NsiI (yellow), XhoI (cyon) NheI (purple). Nucleotides that have been mutated (using synonym codons to avoid amino acid substitutions) are marked in red highlight.

$$N_{27} = 27^N + \sum_{m=1}^{26} (-1)^m \, C_{27-m}^{27} \, (27-m)^N.$$

$$P(N) = \frac{N_{27}}{27^N}$$

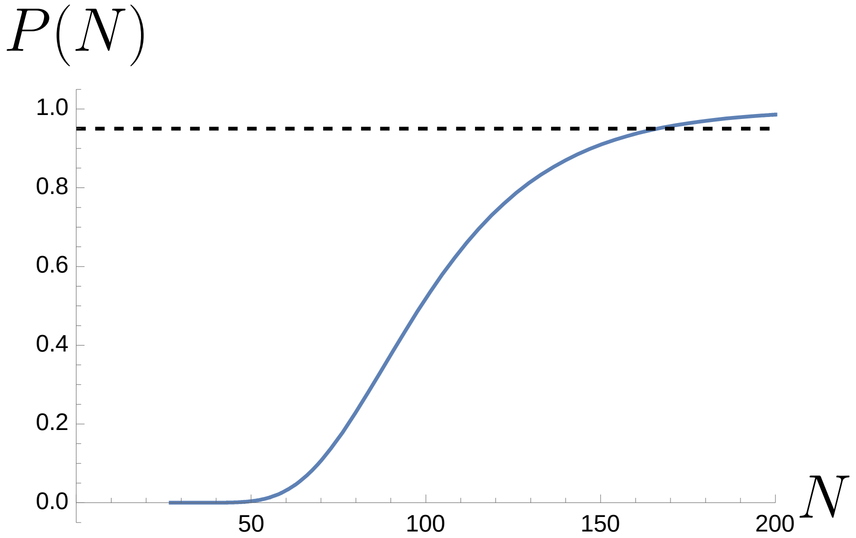

**Supplementary Figure S3:** Estimation of the number of required independent *Synechocystis* transformants (N) in the library to cover the 27 possible RBS-gene combinations in the 3HB operon at sufficienctly high probability P(N).

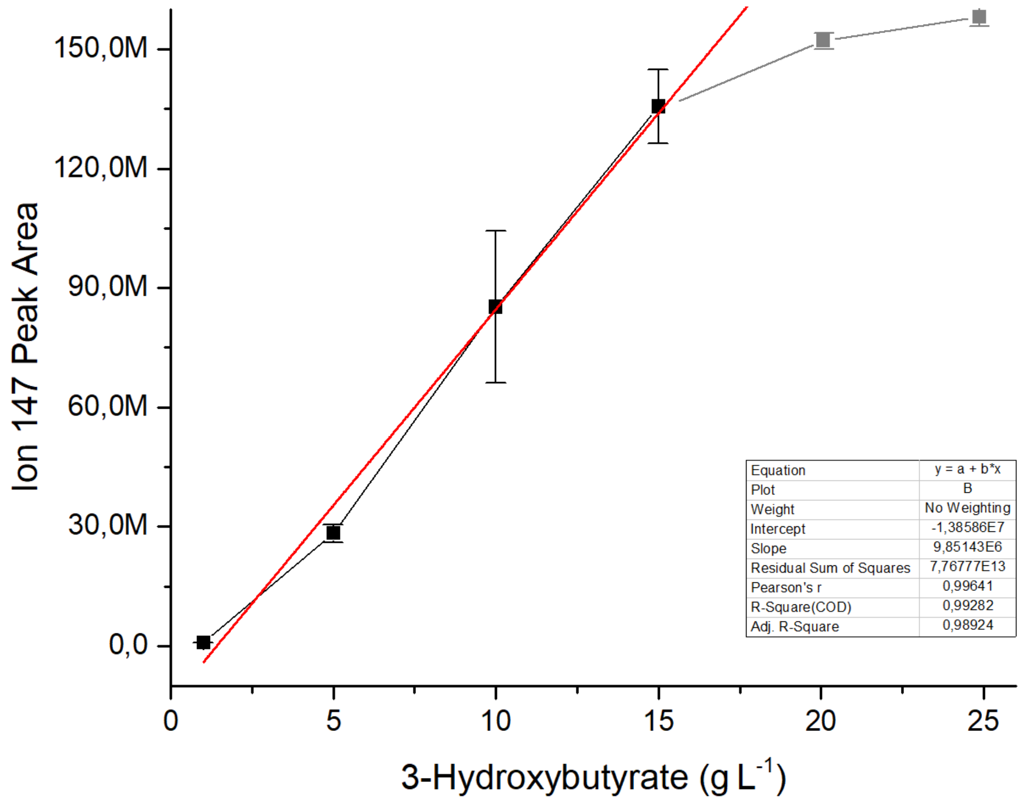

**Supplementary Figure S4**: **3HB GC-MS standard curve** used for the quantitation of 3HB based on the MS ion peak m/z 147. The standard curve has been prepared using known dilutions of a commercial 3HB standard, that have been derivatized with the same method as used for analytical samples. The averages and standard deviations represent three independent replicates (n = 3)

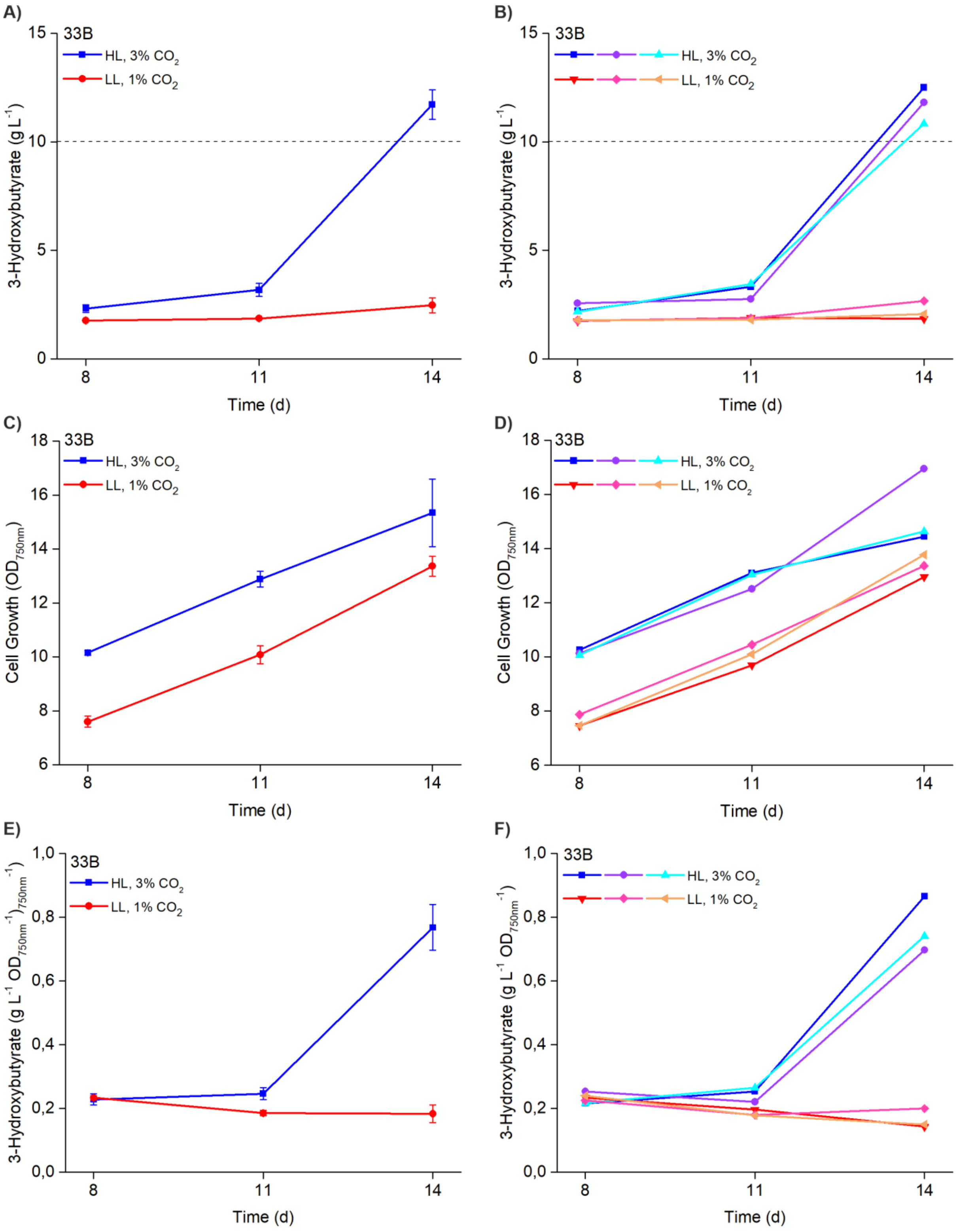

**Supplementary Figure S5**: **Strain 33B complete measured profile for 3HB production and growth** measured under 200 µmol photons $m^{-2}s^{-1}$ continuous light with 3% $CO_2$ (blue profile) and 50 µmol photons $m^{-2}s^{-1}$ continuous light 1% $CO_2$ (red profile) at three different time points (d8, d11, d14) in a two-week batch culture. The data shows **A)** the averaged 3HB accumulation profiles, **B)** the independent 3HB datasets measured from three separate cultivations, **C)** the averaged $OD_{750nm}$ values, **D)** the independent $OD_{750nm}$ values measured from three separate cultivations, **E)** the averaged 3HB concentrations normalized to $OD_{750nm}$, and **F)** the three independent 3HB datasets normalized to $OD_{750nm}$ as measured for each strain in three replicates (n = 3).

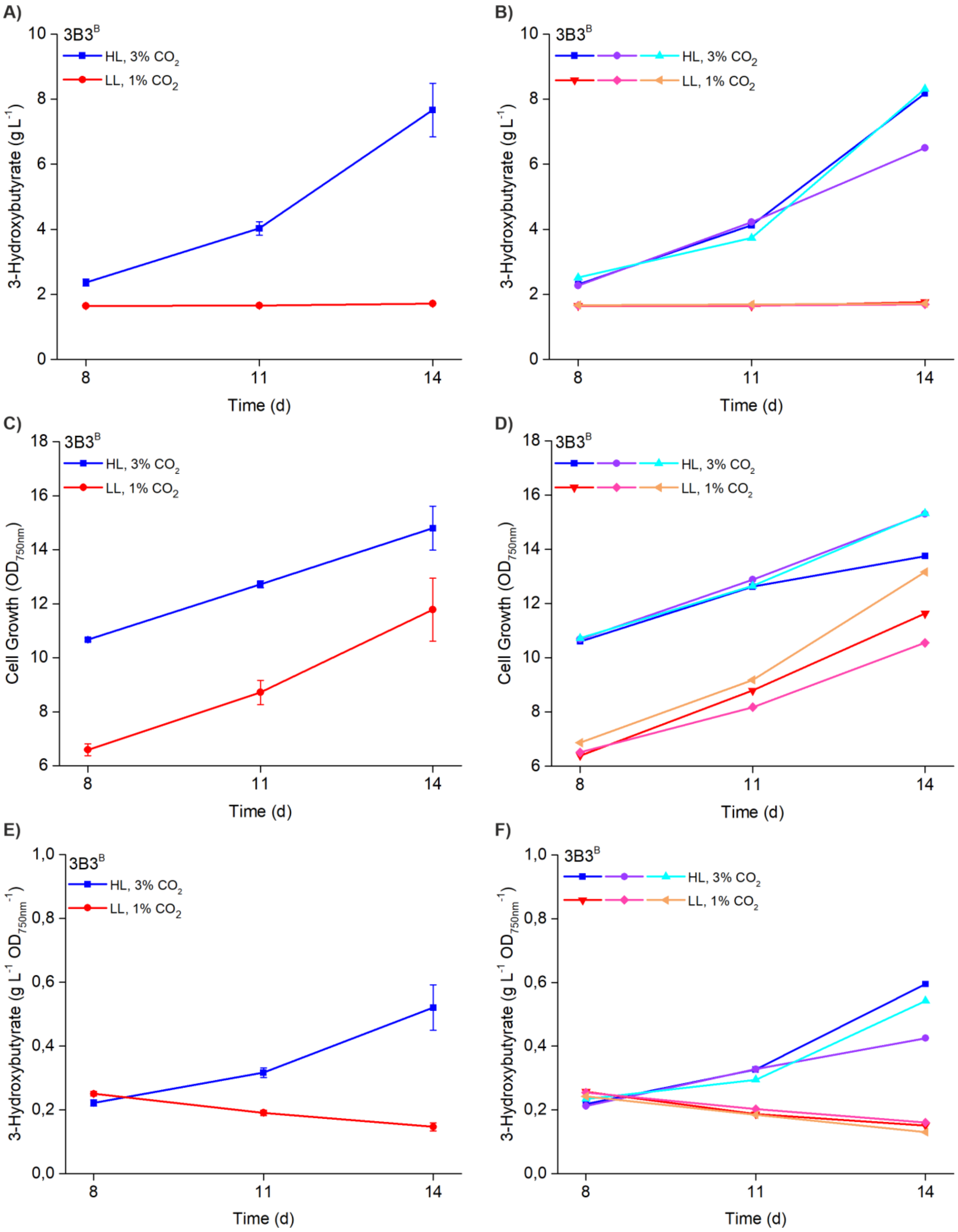

**Supplementary Figure S6: Strain 3B3$^{B}$ complete measured profile for 3HB production and growth** measured under 200 µmol photons m$^{-2}$s$^{-1}$ continuous light with 3% $CO_2$ (blue profile) and 50 µmol photons m$^{-2}$s$^{-1}$ continuous light 1% $CO_2$ (red profile) at three different time points (d8, d11, d14) in a two-week batch culture. The data shows **A)** the averaged 3HB accumulation profiles, **B)** the independent 3HB datasets measured from three separate cultivations, **C)** the averaged $OD_{750nm}$ values, **D)** the independent $OD_{750nm}$ values measured from three separate cultivations, **E)** the averaged 3HB concentrations normalized to $OD_{750nm}$, and **F)** the three independent 3HB datasets normalized to $OD_{750nm}$ as measured for each strain in three replicates (n = 3).

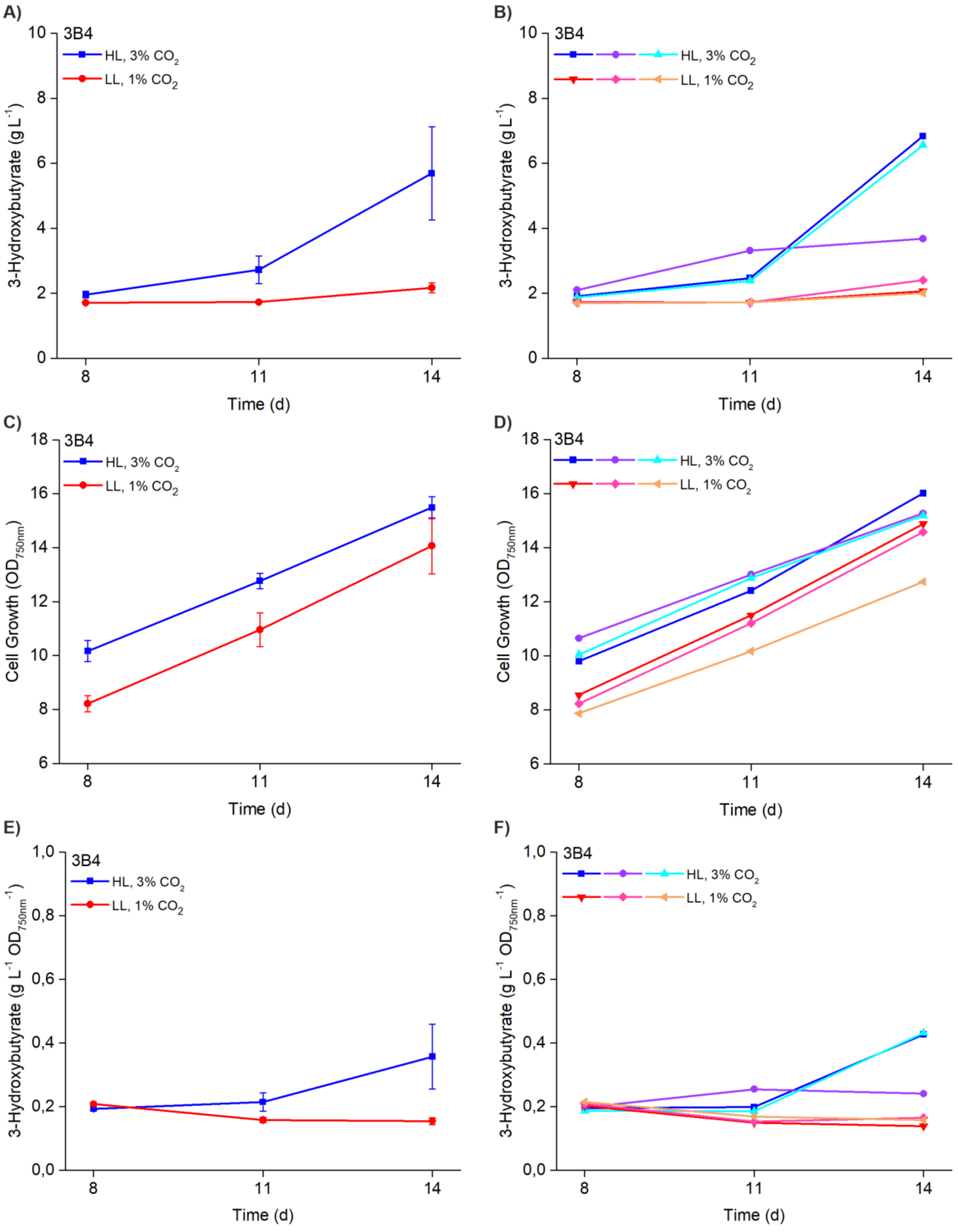

**Supplementary Figure S7: Strain 3B4 complete measured profile for 3HB production and growth** measured under 200 µmol photons $m^{-2}s^{-1}$ continuous light with 3% $CO_2$ (blue profile) and 50 µmol photons $m^{-2}s^{-1}$ continuous light 1% $CO_2$ (red profile) at three different time points (d8, d11, d14) in a two-week batch culture. The data shows **A)** the averaged 3HB accumulation profiles, **B)** the independent 3HB datasets measured from three separate cultivations, **C)** the averaged $OD_{750nm}$ values, **D)** the independent $OD_{750nm}$ values measured from three separate cultivations, **E)** the averaged 3HB concentrations normalized to $OD_{750nm}$, and **F)** the three independent 3HB datasets normalized to $OD_{750nm}$ as measured for each strain in three replicates (n = 3).

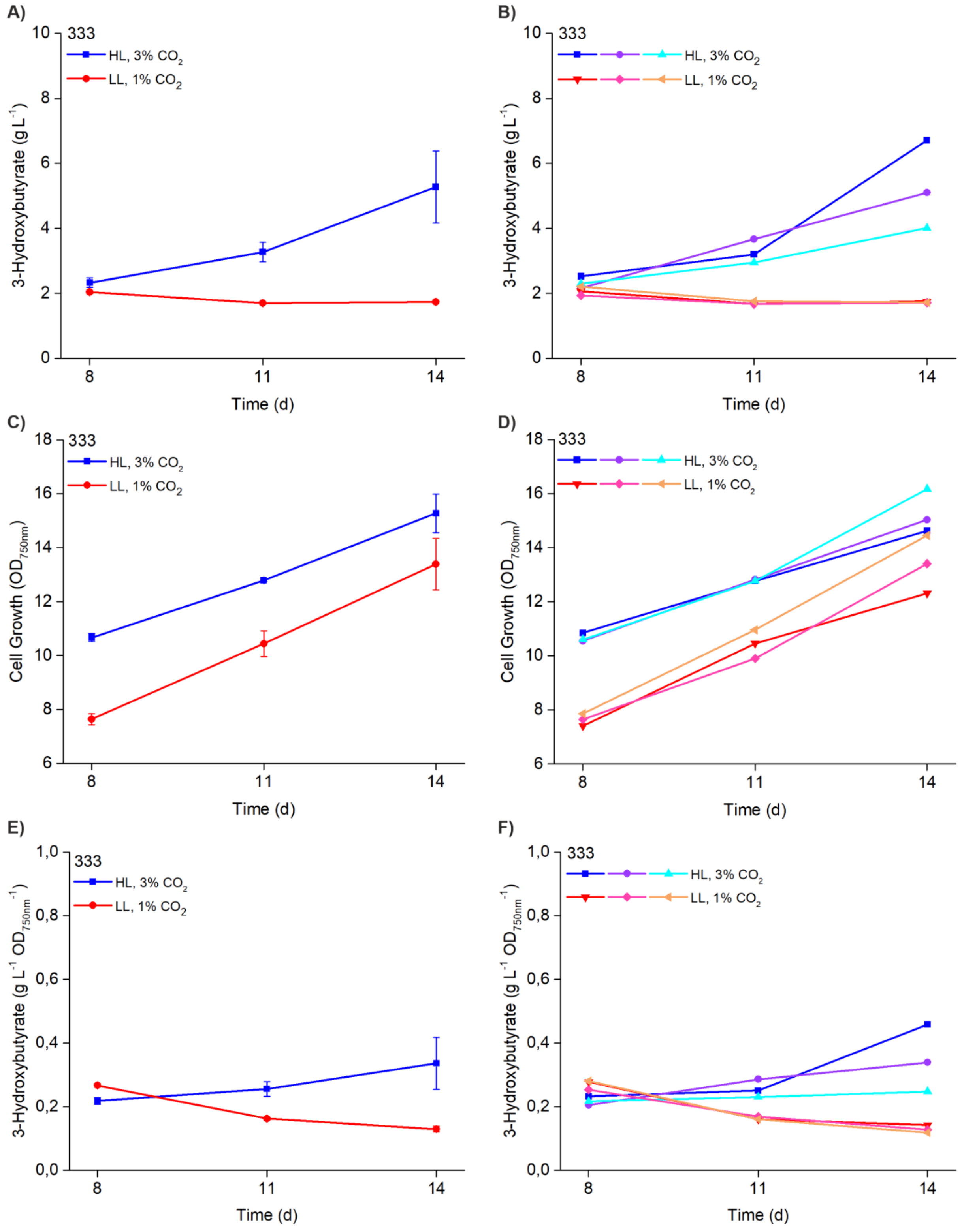

**Supplementary Figure S8: Strain 333 complete measured profile for 3HB production and growth** measured under 200 µmol photons $m^{-2}s^{-1}$ continuous light with 3% $CO_2$ (blue profile) and 50 µmol photons $m^{-2}s^{-1}$ continuous light 1% $CO_2$ (red profile) at three different time points (d8, d11, d14) in a two-week batch culture. The data shows **A)** the averaged 3HB accumulation profiles, **B)** the independent 3HB datasets measured from three separate cultivations, **C)** the averaged $OD_{750nm}$ values, **D)** the independent $OD_{750nm}$ values measured from three separate cultivations, **E)** the averaged 3HB concentrations normalized to $OD_{750nm}$, and **F)** the three independent 3HB datasets normalized to $OD_{750nm}$ as measured for each strain in three replicates (n = 3).

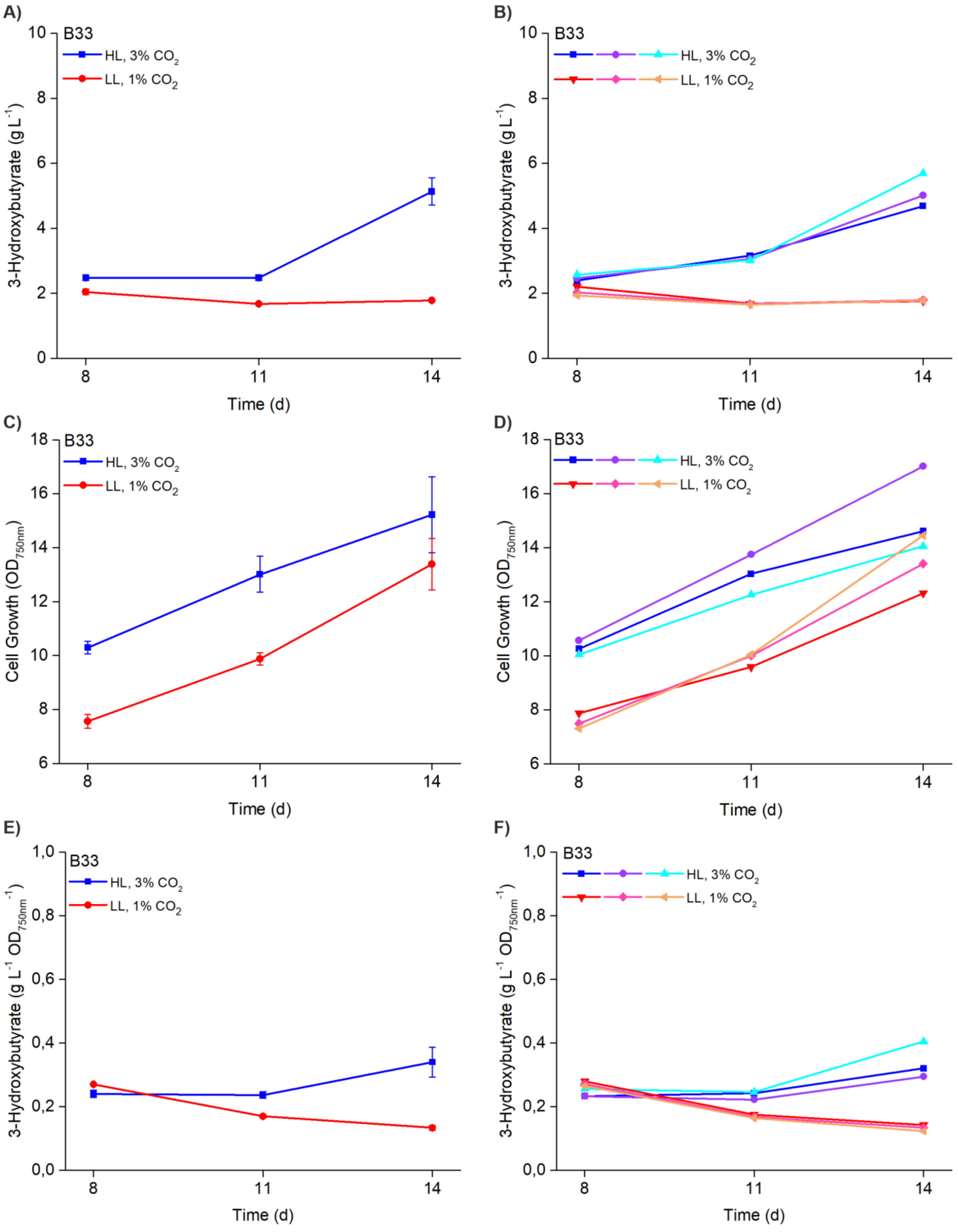

**Supplementary Figure S9: Strain B33 complete measured profile for 3HB production and growth** measured under 200 µmol photons $m^{-2}s^{-1}$ continuous light with 3% $CO_2$ (blue profile) and 50 µmol photons $m^{-2}s^{-1}$ continuous light 1% $CO_2$ (red profile) at three different time points (d8, d11, d14) in a two-week batch culture. The data shows **A)** the averaged 3HB accumulation profiles, **B)** the independent 3HB datasets measured from three separate cultivations, **C)** the averaged $OD_{750nm}$ values, **D)** the independent $OD_{750nm}$ values measured from three separate cultivations, **E)** the averaged 3HB concentrations normalized to $OD_{750nm}$, and **F)** the three independent 3HB datasets normalized to $OD_{750nm}$ as measured for each strain in three replicates (n = 3).

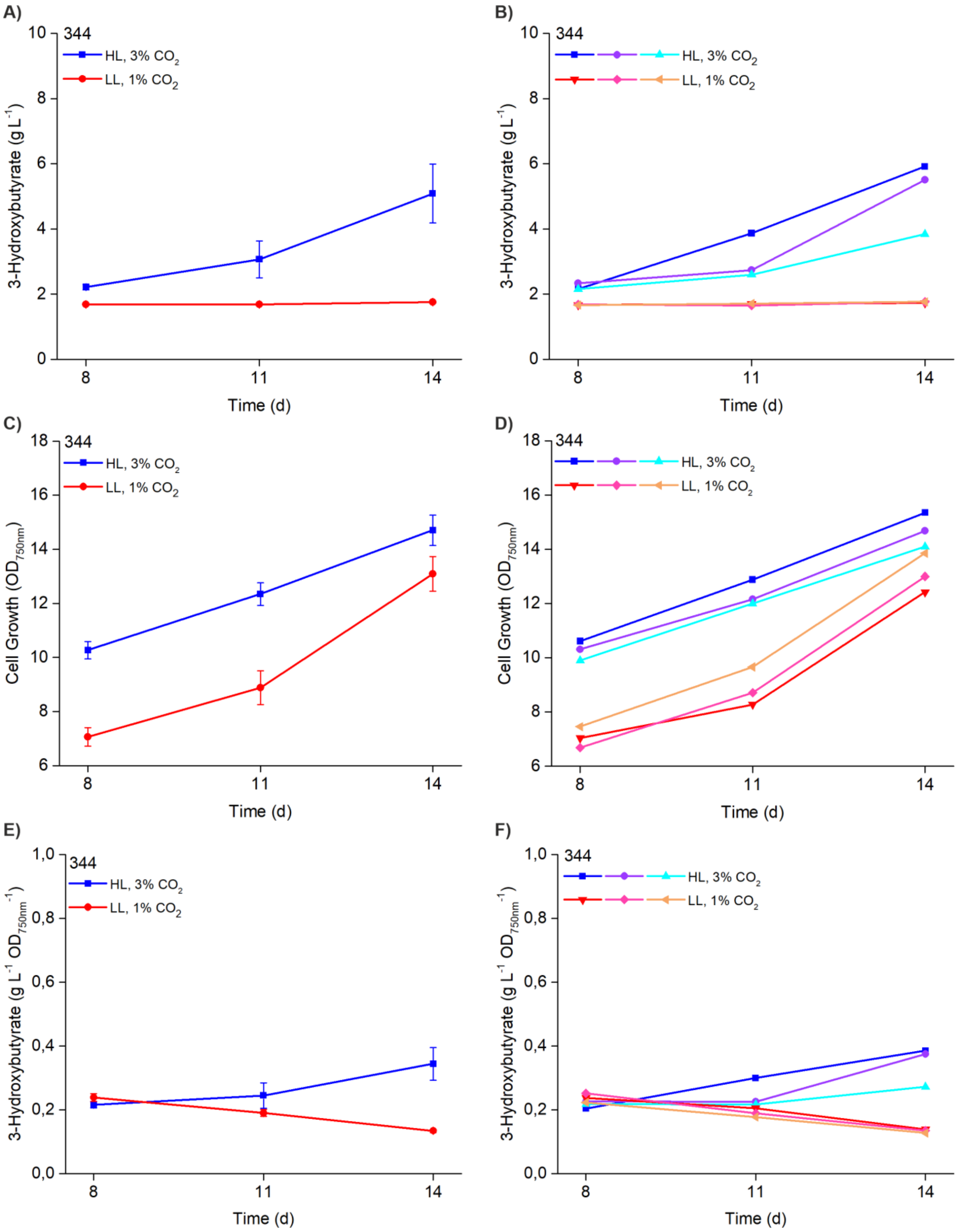

**Supplementary Figure S10: Strain 344 complete measured profile for 3HB production and growth** measured under 200 µmol photons $m^{-2}s^{-1}$ continuous light with 3% $CO_2$ (blue profile) and 50 µmol photons $m^{-2}s^{-1}$ continuous light 1% $CO_2$ (red profile) at three different time points (d8, d11, d14) in a two-week batch culture. The data shows **A)** the averaged 3HB accumulation profiles, **B)** the independent 3HB datasets measured from three separate cultivations, **C)** the averaged $OD_{750nm}$ values, **D)** the independent $OD_{750nm}$ values measured from three separate cultivations, **E)** the averaged 3HB concentrations normalized to $OD_{750nm}$, and **F)** the three independent 3HB datasets normalized to $OD_{750nm}$ as measured for each strain in three replicates (n = 3).

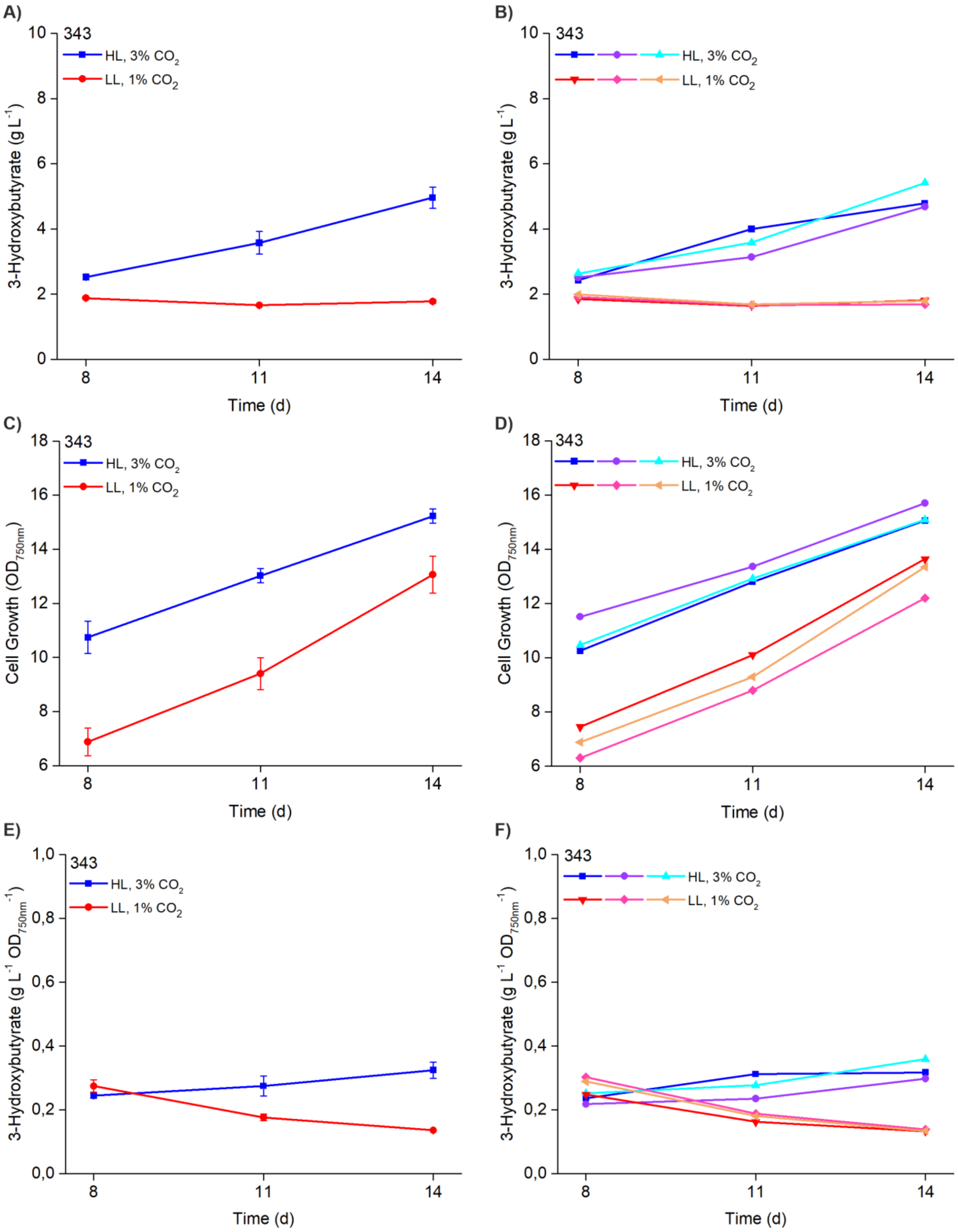

**Supplementary Figure S11: Strain 343 complete measured profile for 3HB production and growth** measured under 200 µmol photons $m^{-2}s^{-1}$ continuous light with 3% $CO_2$ (blue profile) and 50 µmol photons $m^{-2}s^{-1}$ continuous light 1% $CO_2$ (red profile) at three different time points (d8, d11, d14) in a two-week batch culture. The data shows **A)** the averaged 3HB accumulation profiles, **B)** the independent 3HB datasets measured from three separate cultivations, **C)** the averaged $OD_{750nm}$ values, **D)** the independent $OD_{750nm}$ values measured from three separate cultivations, **E)** the averaged 3HB concentrations normalized to $OD_{750nm}$, and **F)** the three independent 3HB datasets normalized to $OD_{750nm}$ as measured for each strain in three replicates (n = 3).

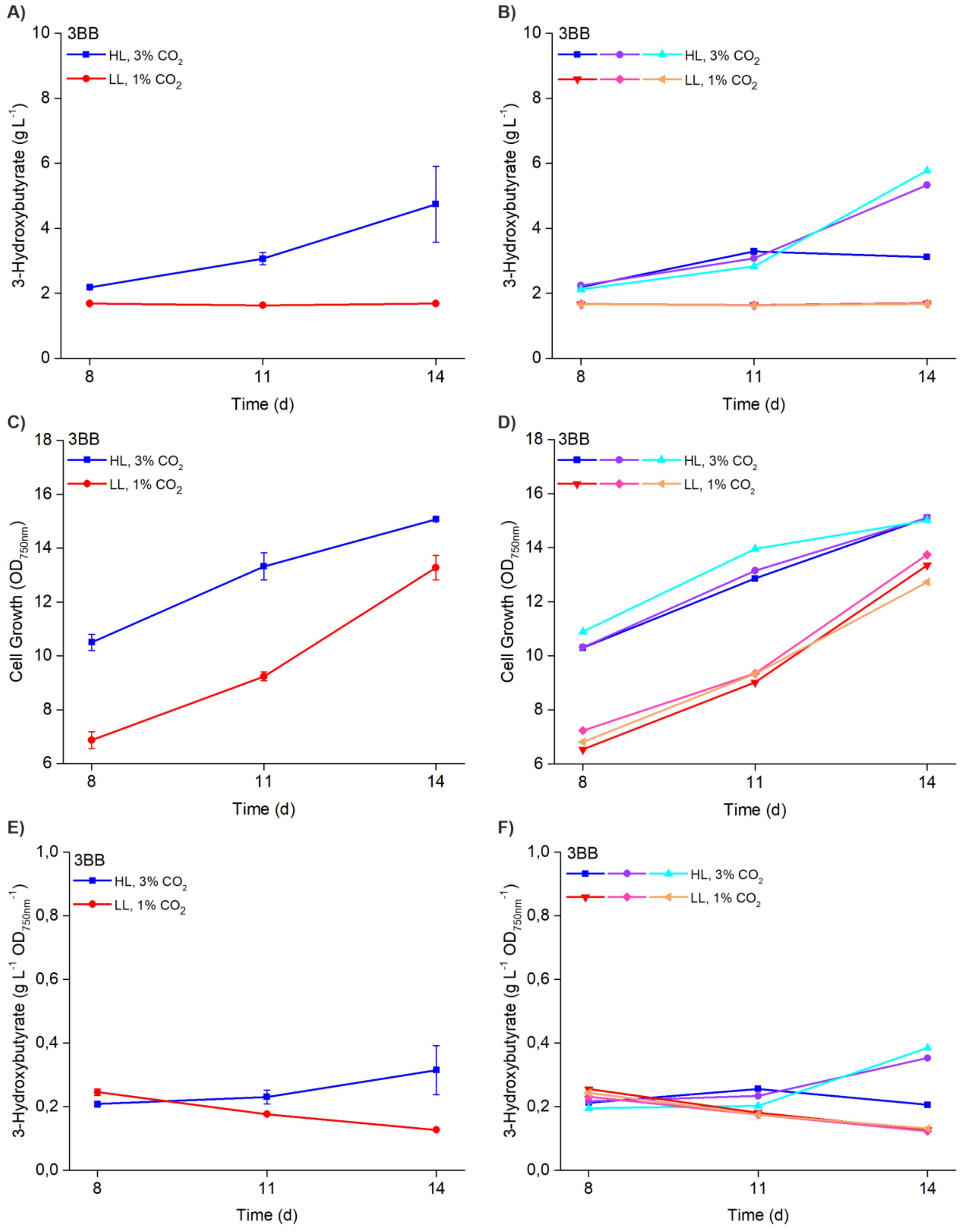

**Supplementary Figure S12: Strain 3BB complete measured profile for 3HB production and growth** measured under 200 µmol photons $m^{-2}s^{-1}$ continuous light with 3% $CO_2$ (blue profile) and 50 µmol photons $m^{-2}s^{-1}$ continuous light 1% $CO_2$ (red profile) at three different time points (d8, d11, d14) in a two-week batch culture. The data shows **A)** the averaged 3HB accumulation profiles, **B)** the independent 3HB datasets measured from three separate cultivations, **C)** the averaged $OD_{750nm}$ values, **D)** the independent $OD_{750nm}$ values measured from three separate cultivations, **E)** the averaged 3HB concentrations normalized to $OD_{750nm}$, and **F)** the three independent 3HB datasets normalized to $OD_{750nm}$ as measured for each strain in three replicates (n = 3).

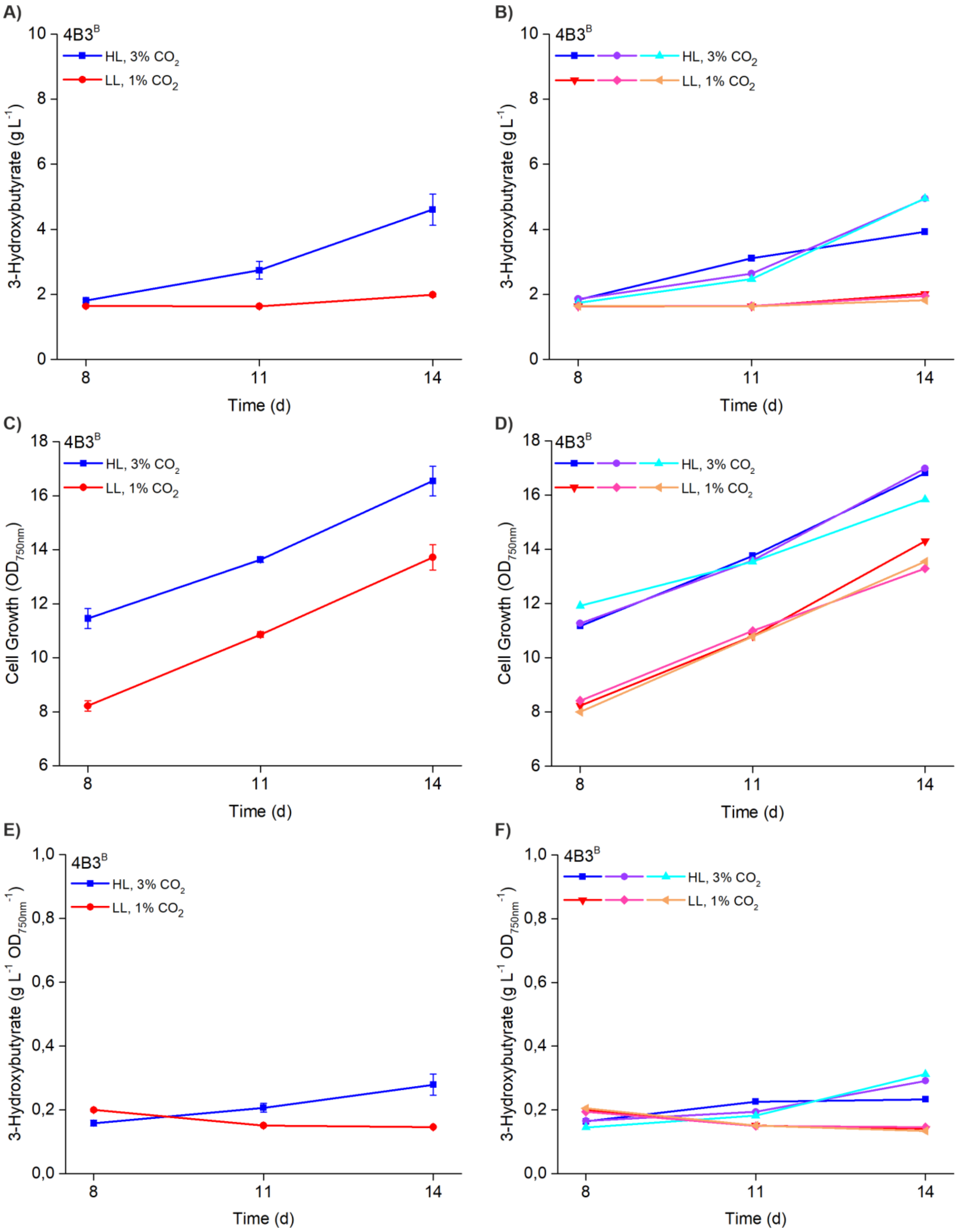

**Supplementary Figure S13: Strain 4B3$^B$ complete measured profile for 3HB production and growth** measured under 200 µmol photons $m^{-2}s^{-1}$ continuous light with 3% $CO_2$ (blue profile) and 50 µmol photons $m^{-2}s^{-1}$ continuous light 1% $CO_2$ (red profile) at three different time points (d8, d11, d14) in a two-week batch culture. The data shows **A)** the averaged 3HB accumulation profiles, **B)** the independent 3HB datasets measured from three separate cultivations, **C)** the averaged $OD_{750nm}$ values, **D)** the independent $OD_{750nm}$ values measured from three separate cultivations, **E)** the averaged 3HB concentrations normalized to $OD_{750nm}$, and **F)** the three independent 3HB datasets normalized to $OD_{750nm}$ as measured for each strain in three replicates (n = 3).

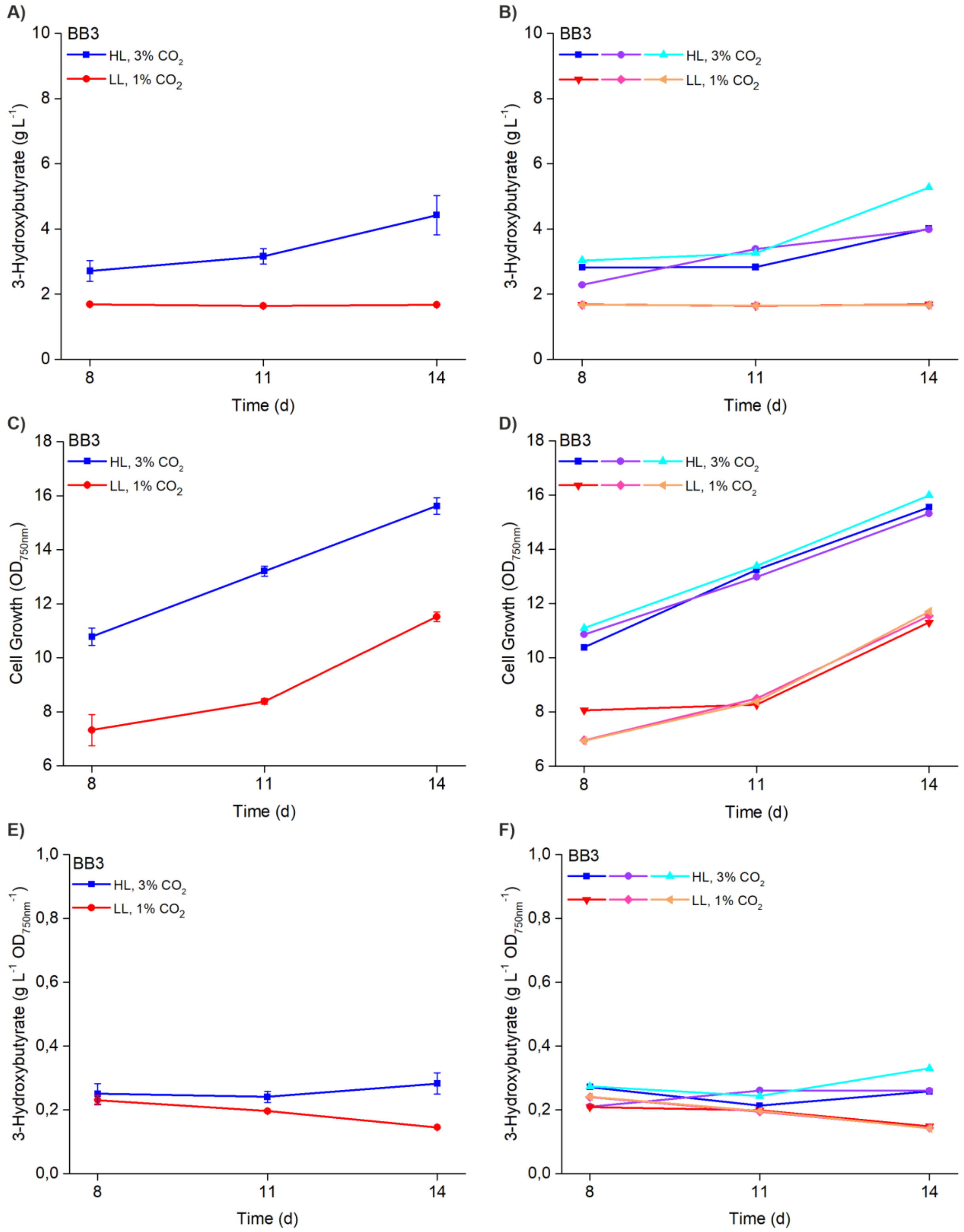

**Supplementary Figure S14: Strain BB3 complete measured profile for 3HB production and growth** measured under 200 µmol photons $m^{-2}s^{-1}$ continuous light with 3% $CO_2$ (blue profile) and 50 µmol photons $m^{-2}s^{-1}$ continuous light 1% $CO_2$ (red profile) at three different time points (d8, d11, d14) in a two-week batch culture. The data shows **A)** the averaged 3HB accumulation profiles, **B)** the independent 3HB datasets measured from three separate cultivations, **C)** the averaged $OD_{750nm}$ values, **D)** the independent $OD_{750nm}$ values measured from three separate cultivations, **E)** the averaged 3HB concentrations normalized to $OD_{750nm}$, and **F)** the three independent 3HB datasets normalized to $OD_{750nm}$ as measured for each strain in three replicates (n = 3).

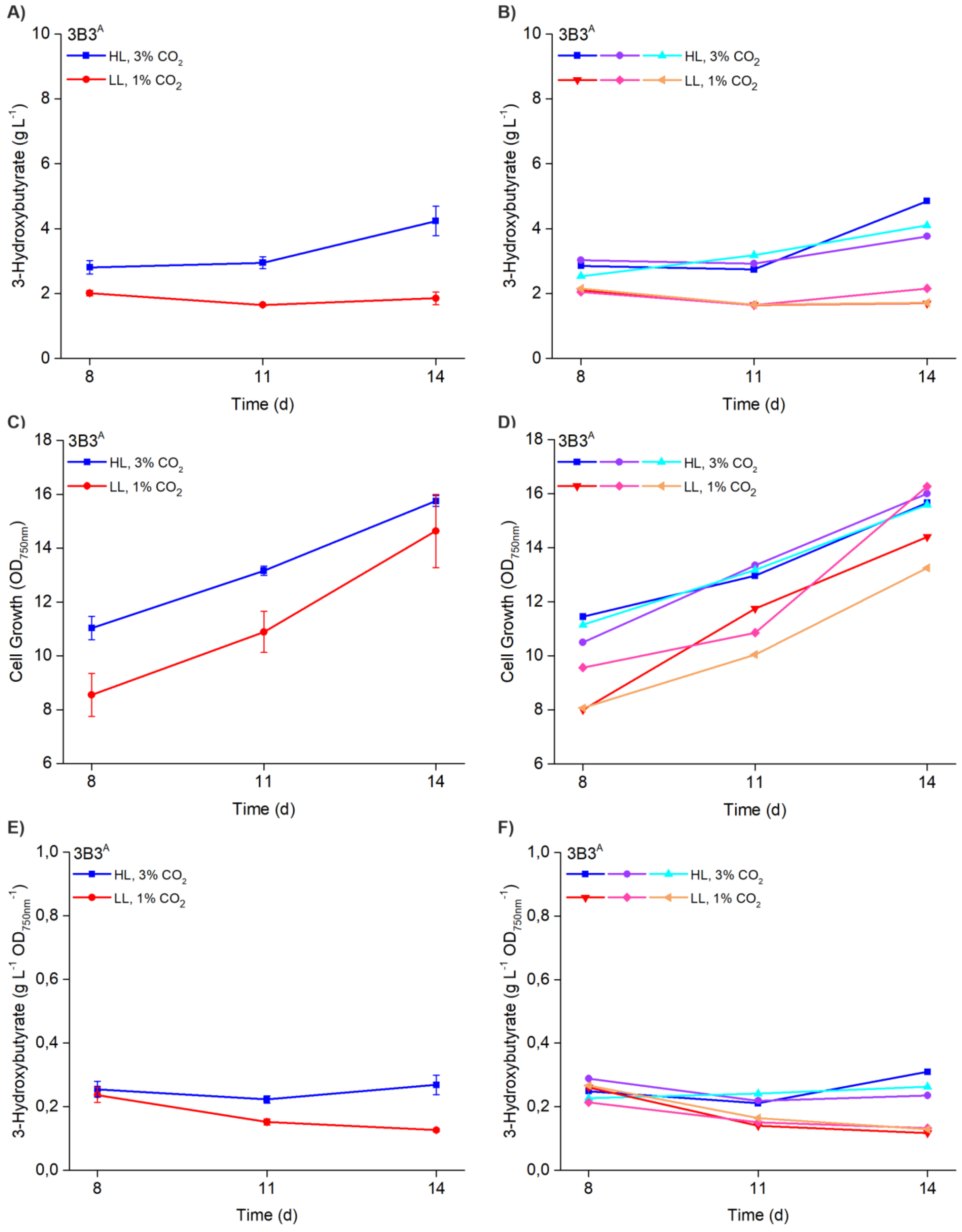

**Supplementary Figure S15: Strain 3B3$^{A}$ complete measured profile for 3HB production and growth** measured under 200 µmol photons $m^{-2}s^{-1}$ continuous light with 3% $CO_2$ (blue profile) and 50 µmol photons $m^{-2}s^{-1}$ continuous light 1% $CO_2$ (red profile) at three different time points (d8, d11, d14) in a two-week batch culture. The data shows **A)** the averaged 3HB accumulation profiles, **B)** the independent 3HB datasets measured from three separate cultivations, **C)** the averaged $OD_{750nm}$ values, **D)** the independent $OD_{750nm}$ values measured from three separate cultivations, **E)** the averaged 3HB concentrations normalized to $OD_{750nm}$, and **F)** the three independent 3HB datasets normalized to $OD_{750nm}$ as measured for each strain in three replicates (n = 3).

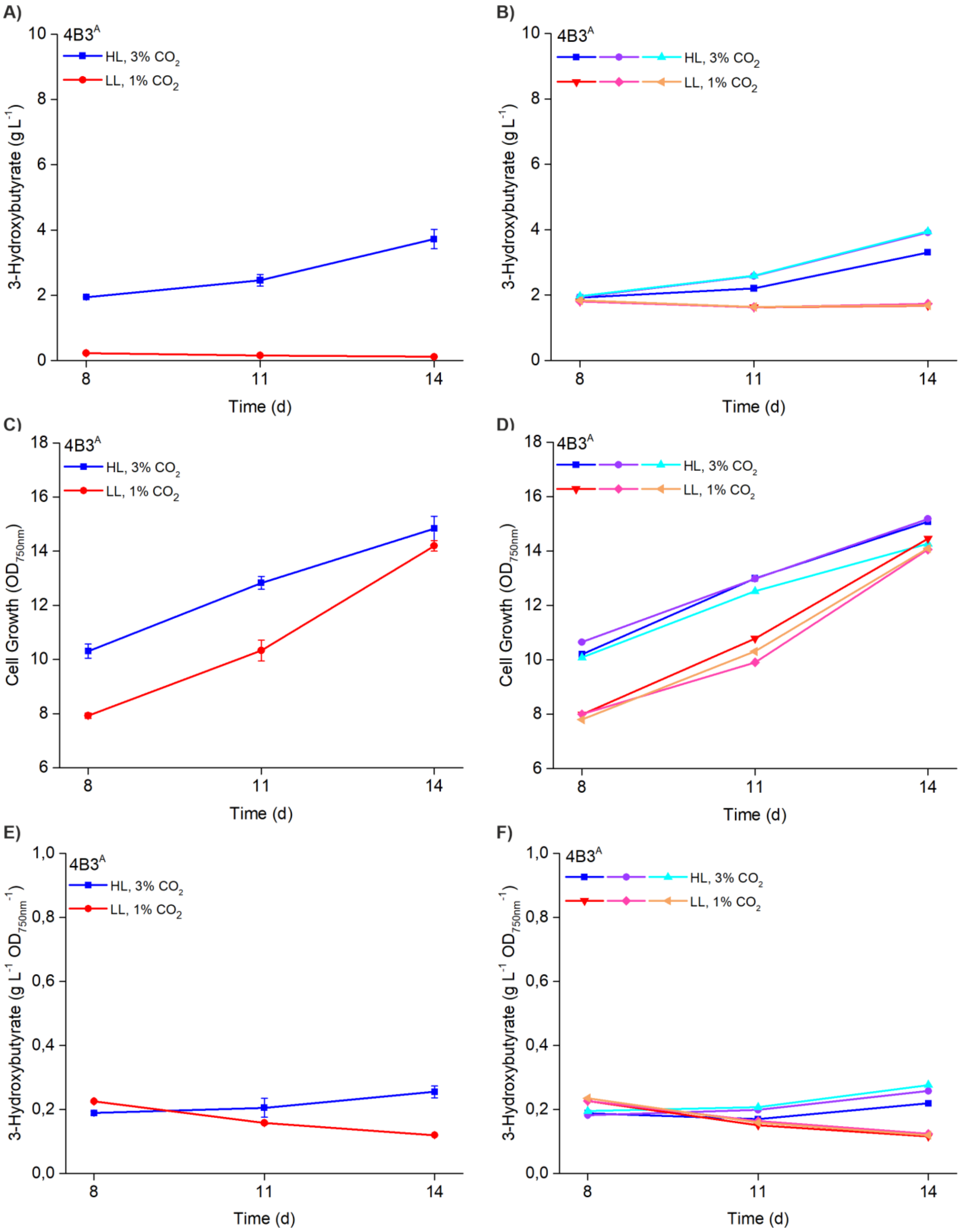

**Supplementary Figure S16: Strain 4B3$^{A}$ complete measured profile for 3HB production and growth** measured under 200 µmol photons $m^{-2}s^{-1}$ continuous light with 3% $CO_2$ (blue profile) and 50 µmol photons $m^{-2}s^{-1}$ continuous light 1% $CO_2$ (red profile) at three different time points (d8, d11, d14) in a two-week batch culture. The data shows **A)** the averaged 3HB accumulation profiles, **B)** the independent 3HB datasets measured from three separate cultivations, **C)** the averaged $OD_{750nm}$ values, **D)** the independent $OD_{750nm}$ values measured from three separate cultivations, **E)** the averaged 3HB concentrations normalized to $OD_{750nm}$, and **F)** the three independent 3HB datasets normalized to $OD_{750nm}$ as measured for each strain in three replicates (n = 3).

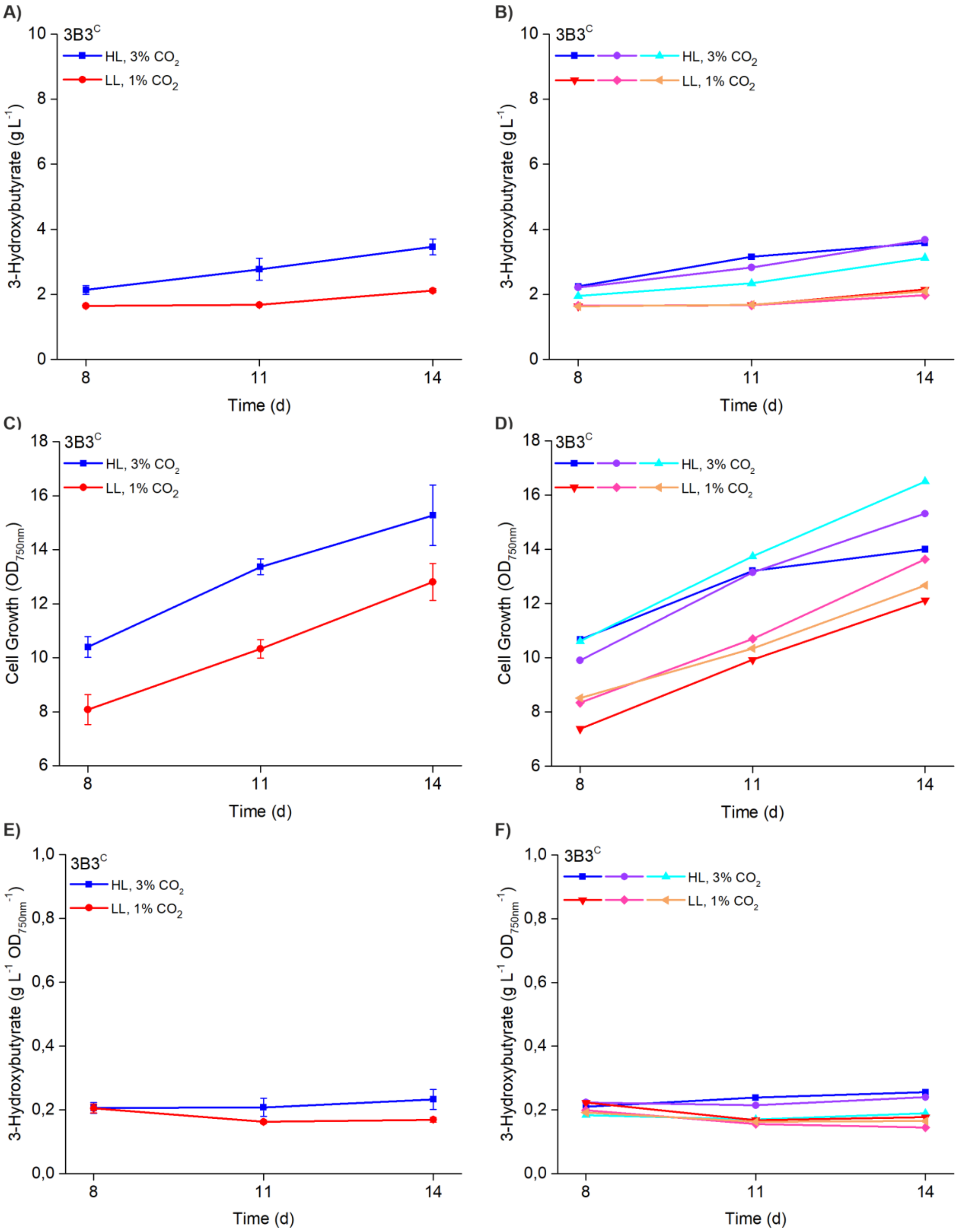

**Supplementary Figure S17: Strain 3B3$^C$ complete measured profile for 3HB production and growth** measured under 200 µmol photons $m^{-2}s^{-1}$ continuous light with 3% $CO_2$ (blue profile) and 50 µmol photons $m^{-2}s^{-1}$ continuous light 1% $CO_2$ (red profile) at three different time points (d8, d11, d14) in a two-week batch culture. The data shows **A)** the averaged 3HB accumulation profiles, **B)** the independent 3HB datasets measured from three separate cultivations, **C)** the averaged $OD_{750nm}$ values, **D)** the independent $OD_{750nm}$ values measured from three separate cultivations, **E)** the averaged 3HB concentrations normalized to $OD_{750nm}$, and **F)** the three independent 3HB datasets normalized to $OD_{750nm}$ as measured for each strain in three replicates (n = 3).

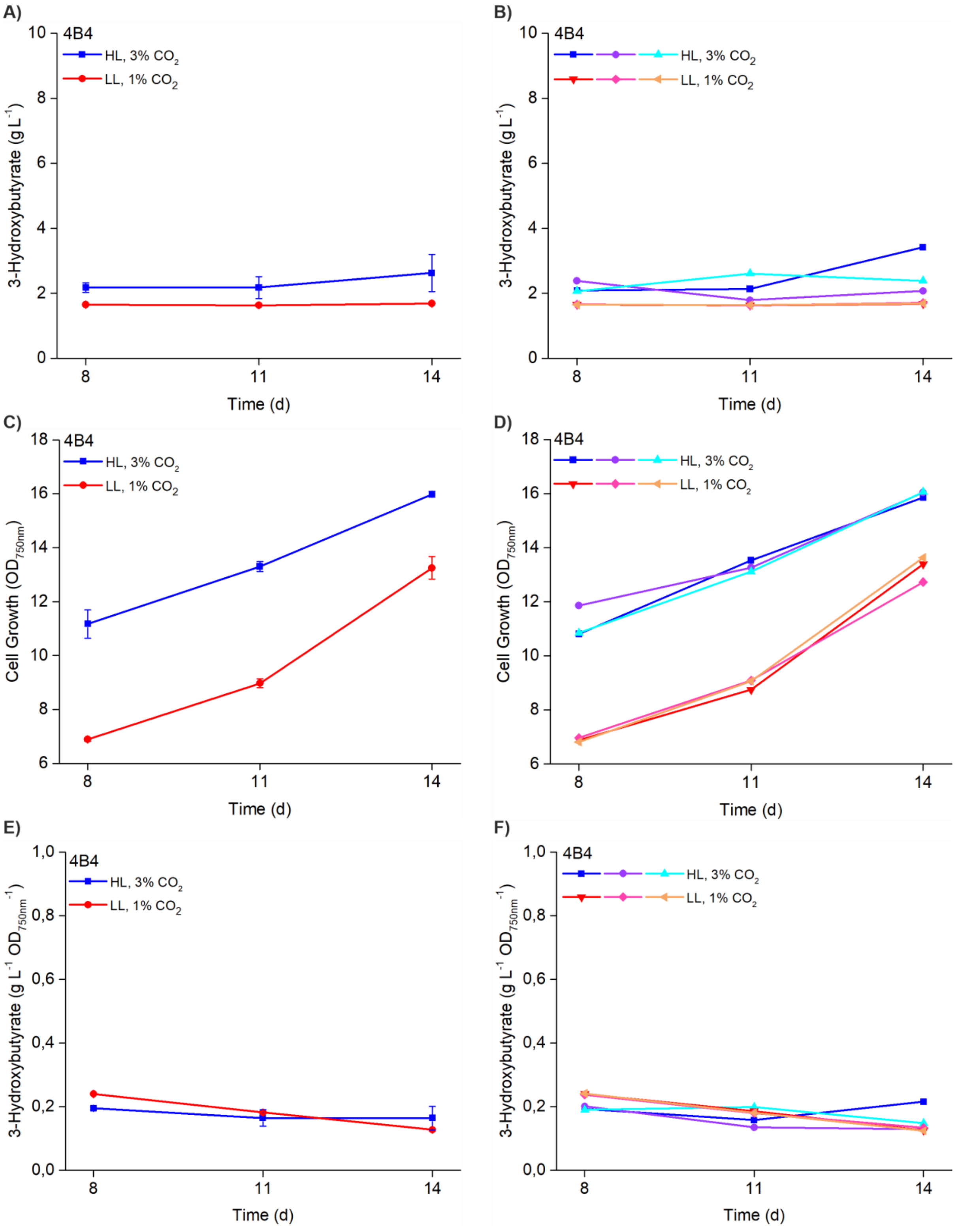

**Supplementary Figure S18**: Strain 4B4 **complete measured profile for 3HB production and growth** measured under 200 µmol photons $m^{-2}s^{-1}$ continuous light with 3% $CO_2$ (blue profile) and 50 µmol photons $m^{-2}s^{-1}$ continuous light 1% $CO_2$ (red profile) at three different time points (d8, d11, d14) in a two-week batch culture. The data shows **A)** the averaged 3HB accumulation profiles, **B)** the independent 3HB datasets measured from three separate cultivations, **C)** the averaged $OD_{750nm}$ values, **D)** the independent $OD_{750nm}$ values measured from three separate cultivations, **E)** the averaged 3HB concentrations normalized to $OD_{750nm}$, and **F)** the three independent 3HB datasets normalized to $OD_{750nm}$ as measured for each strain in three replicates (n = 3).

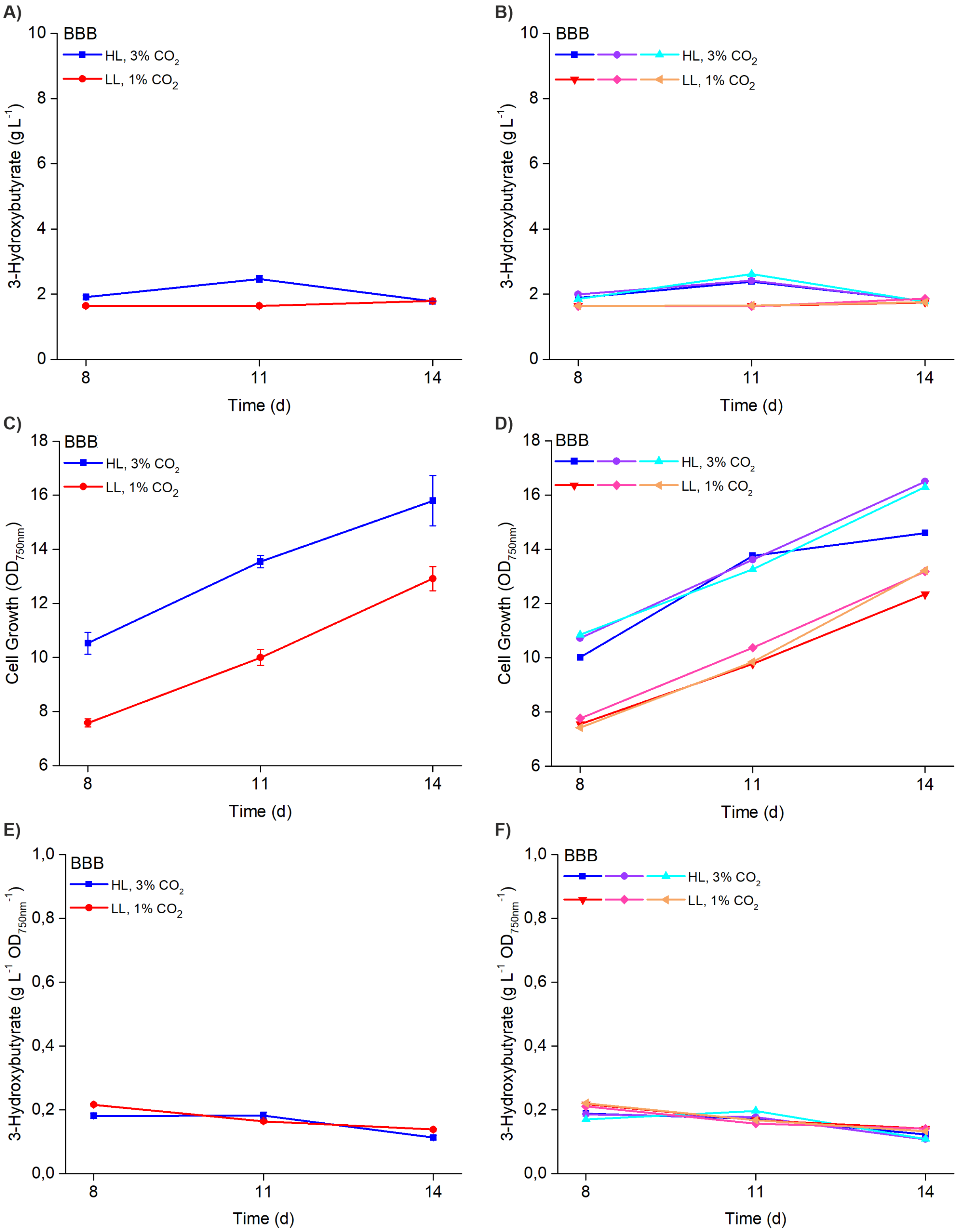

**Supplementary Figure S19: Strain BBB complete measured profile for 3HB production and growth** measured under 200 µmol photons $m^{-2}s^{-1}$ continuous light with 3% $CO_2$ (blue profile) and 50 µmol photons $m^{-2}s^{-1}$ continuous light 1% $CO_2$ (red profile) at three different time points (d8, d11, d14) in a two-week batch culture. The data shows **A)** the averaged 3HB accumulation profiles, **B)** the independent 3HB datasets measured from three separate cultivations, **C)** the averaged $OD_{750nm}$ values, **D)** the independent $OD_{750nm}$ values measured from three separate cultivations, **E)** the averaged 3HB concentrations normalized to $OD_{750nm}$, and **F)** the three independent 3HB datasets normalized to $OD_{750nm}$ as measured for each strain in three replicates (n = 3).

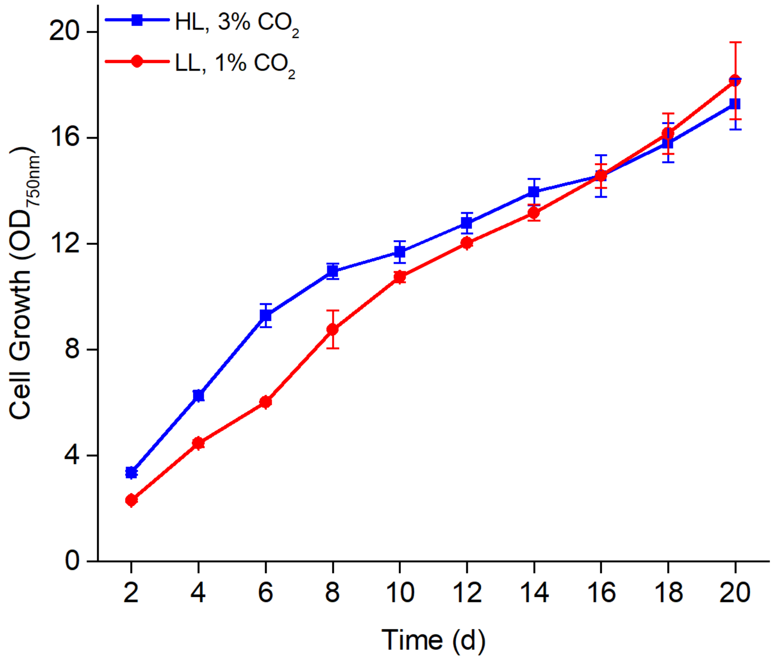

**Supplementary Figure S20: A representative growth OD750mn profile measured every second day over a 20d batch culture** under 200 µmol photons m$^{-2}$s$^{-1}$ continuous light with 3% $CO_2$ (blue profile) and 50 µmol photons m$^{-2}$s$^{-1}$ continuous light 1% $CO_2$ (red profile) for the strain 3B3$^{A}$. The averages and standard deviations represent measurements conducted from three independent cultivations (n = 3).

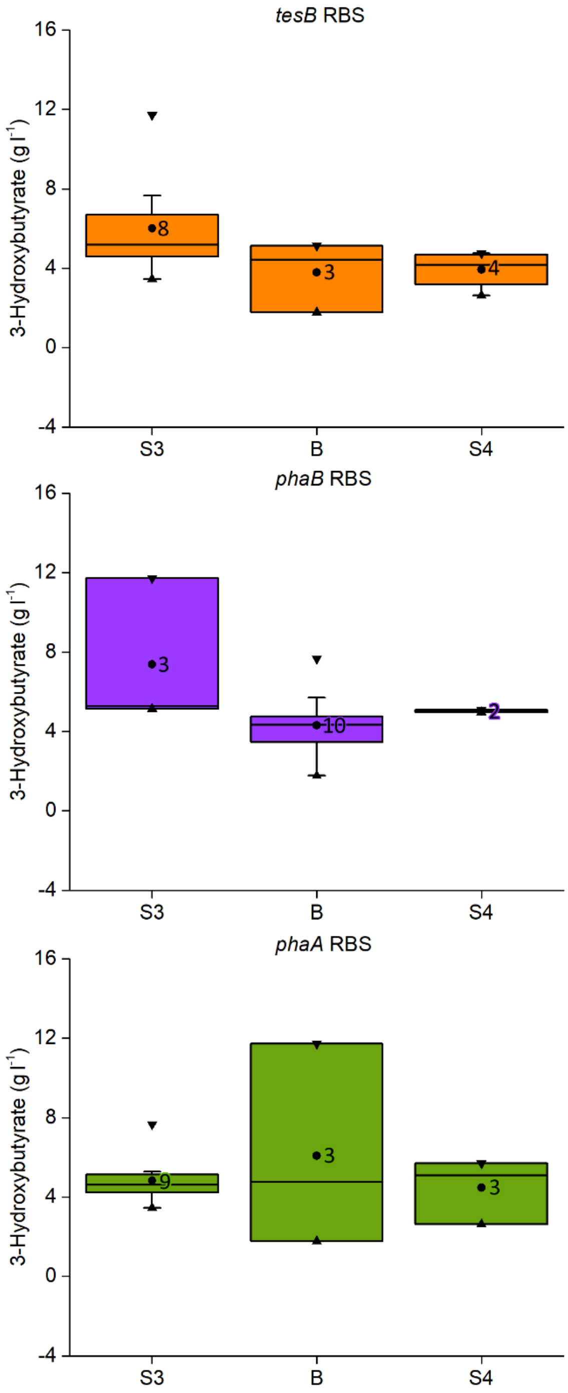

**Supplementary Figure S21**. **Box and whiskers plot illustrating the statistical distribution of measured 3HB level in relation to each gene-RBS pair based on the 15 selected strains**. The box represents the interquartile range (IQR), the line within the box marks the median, the circle within the box marks the average, the whiskers extend from the box to the minimum and maximum values (excluding outliers), and the triangles pointing towards the box mark the minimum and maximum values within the dataset. The N value of each gene-RBS pair in the dataset is indicated next to the average point. Notably, the data does not represent all the possible 27 RBS-gene combinations and is heavily biased by the nonrandom strain selection.

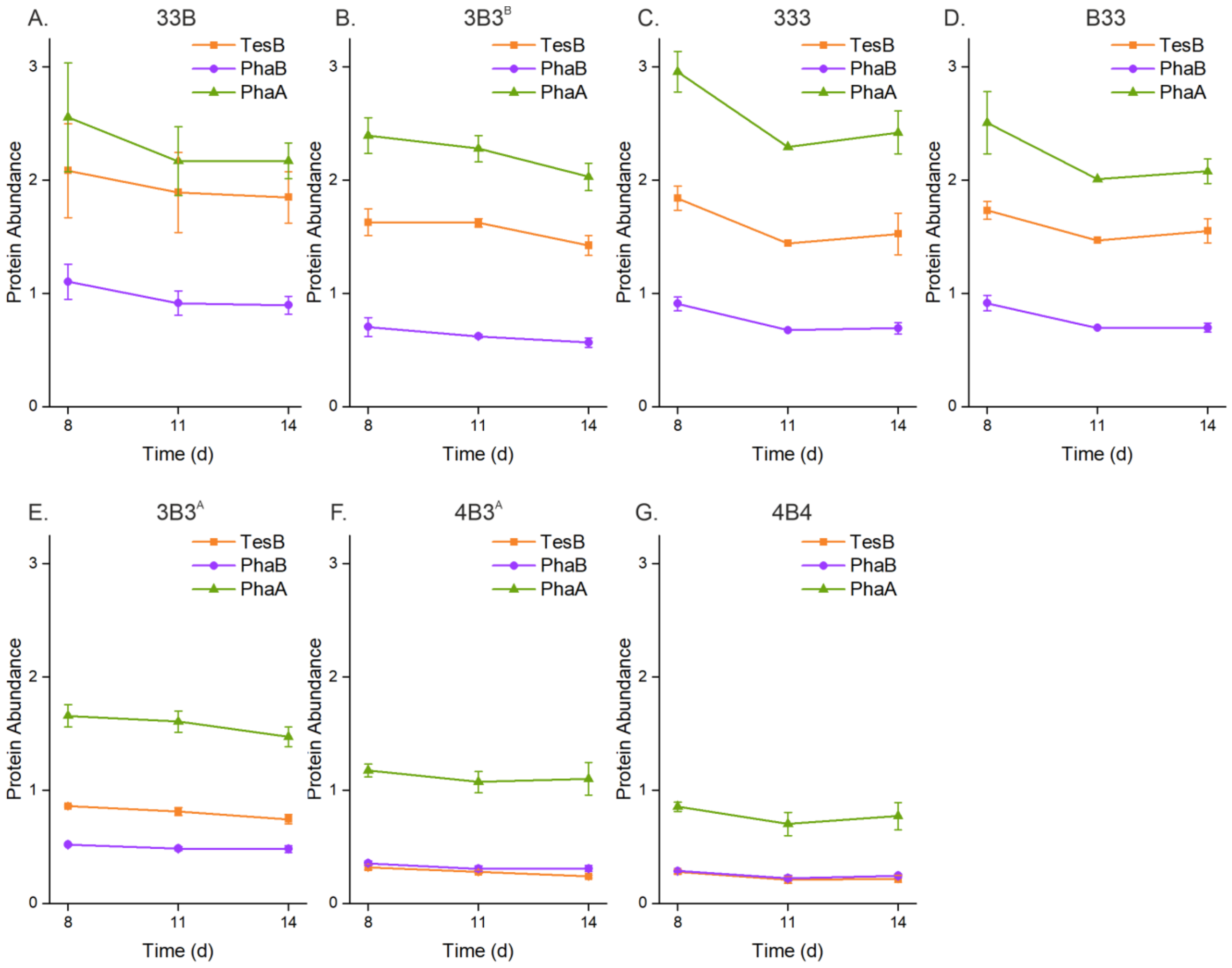

**Supplementary Figure S22: The relative abundance of the 3HB pathway proteins TesB, PhaB and PhaA at three consecutive sampling time points** (d8, d11 and d14) after induction in the seven selected strains **A)** 33B, **B)** $3B3^{B}$, **C)** 333, **D)** B33, **E)** $3B3^{A}$, **F)** $4B3^{A}$ and **G)** 4B4**.** The averages and standard deviations represent measurements conducted from three independent cultivations, based on at least three quantitated peptides per protein (n = 3).

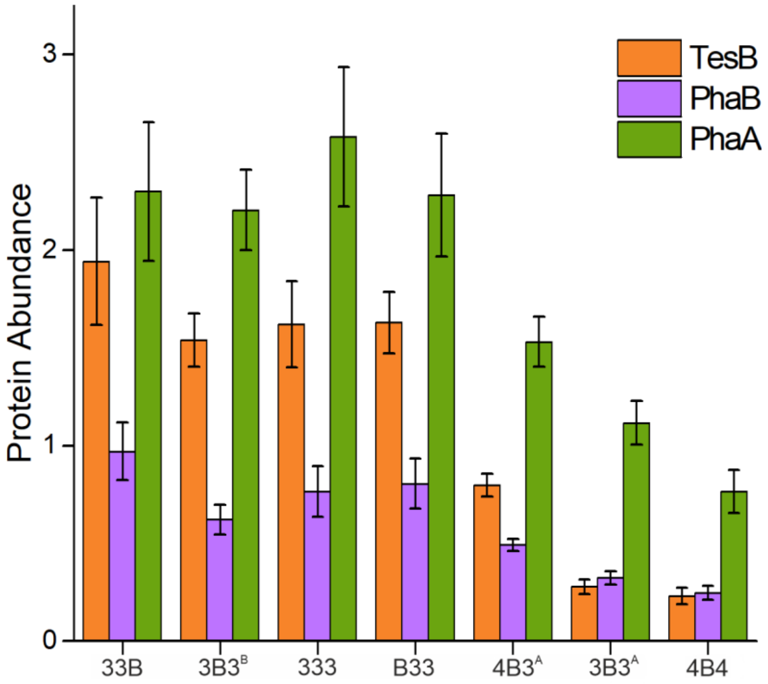

**Supplementary Figure S23. The combined relative abundance of the 3HB pathway proteins TesB, PhaB and PhaA in the seven selected strains.** The bars and standard deviations have been calculated form the combined total data from the three sampling time points d8, d11 and d14 shown in **Supplementary Figure S22** (n = 9).

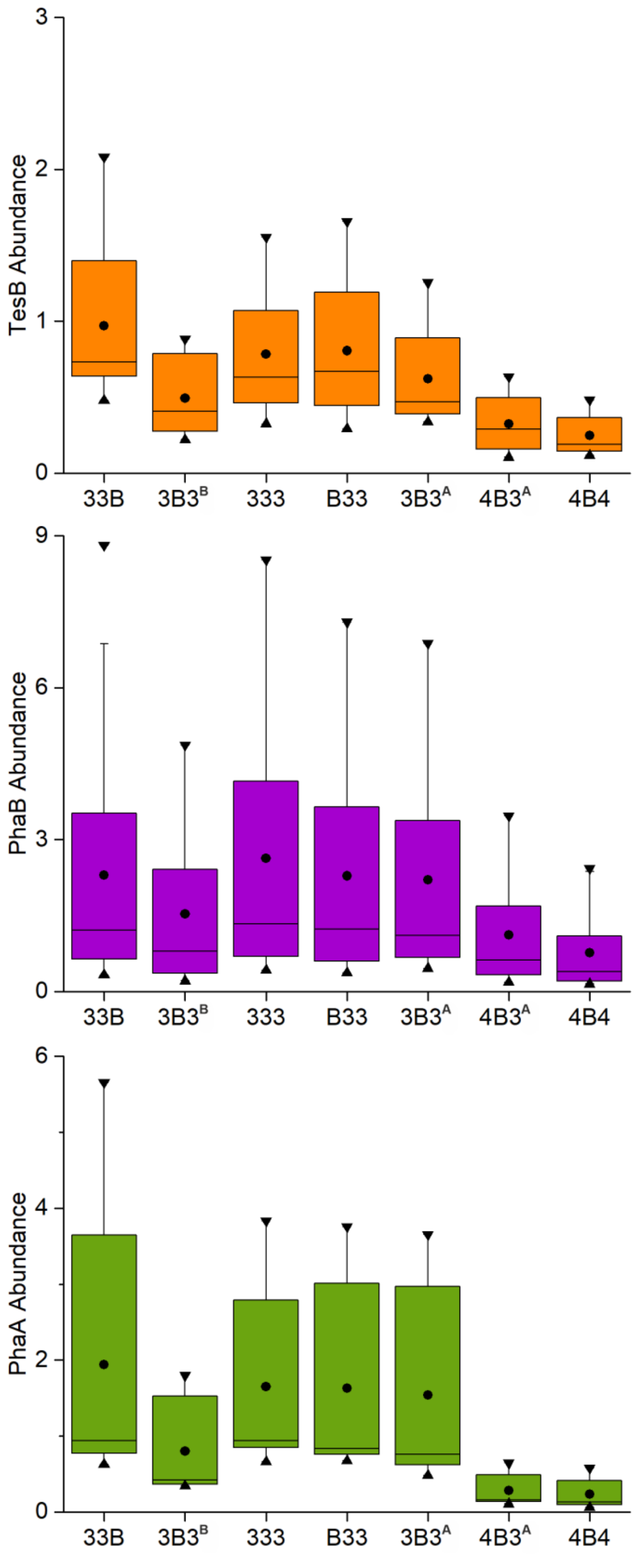

**Supplementary Figure S24. Box and whiskers plot illustrating the abundance of the 3HB pathway proteins TesB, PhaA and PhaB in the seven selected strains** The box represents the interquartile range (IQR), the line within the box marks the median, the circle within the box marks the average, the whiskers extend from the box to the minimum and maximum values (excluding outliers), and the triangles pointing towards the box mark the minimum and maximum values within the dataset.

**Supplementary Table S2. Mann-Whitney U test comparing protein abundance of TesB** in strain 33B against strains 333, 3B3[B], 3B3[A], 4B3, 4B4[A], and B33, showing sample sizes (N), mean abundances, U statistics, p-values, and significance ($p < 0.05$ indicates a significant difference in distributions).

| Comparator Strain | N (33B) | N (Other) | Mean (33B) | Mean (Other) | U Statistic | p-value | Significance |
|---|---|---|---|---|---|---|---|
| 333 | 27 | 27 | 1.285 | 2.156 | 232.0 | 0.0224 | Yes |
| 3B3[B] | 27 | 24 | 1.285 | 1.416 | 386.0 | 0.2458 | No |
| 3B3[A] | 27 | 24 | 1.285 | 0.673 | 523.0 | 0.0002 | Yes |
| 4B3[A] | 27 | 21 | 1.285 | 0.212 | 554.0 | 0.0000 | Yes |
| 4B4 | 27 | 21 | 1.285 | 0.196 | 559.0 | 0.0000 | Yes |
| B33 | 27 | 18 | 1.285 | 1.629 | 220.0 | 0.6022 | No |

**Supplementary Table S3. Mann-Whitney U test comparing protein abundance of PhaA** in strain 33B against strains 333, 3B3$^{B}$, 3B3$^{A}$, 4B3, 4B4$^{A}$, and B33, showing sample sizes (N), mean abundances, U statistics, p-values, and significance (p < 0.05 indicates a significant difference in distributions).

| Comparator Strain | N (33B) | N (Other) | Mean (33B) | Mean (Other) | U Statistic | p-value | Significance |
|---|---|---|---|---|---|---|---|
| 333 | 36 | 36 | 2.262 | 2.134 | 660.0 | 0.8969 | No |
| 3B3$^{B}$ | 36 | 32 | 2.262 | 1.962 | 607.0 | 0.7078 | No |
| 3B3$^{A}$ | 36 | 32 | 2.262 | 1.523 | 698.0 | 0.1355 | No |
| 4B3$^{A}$ | 36 | 28 | 2.262 | 1.366 | 553.0 | 0.5116 | No |
| 4B4 | 36 | 28 | 2.262 | 0.850 | 676.0 | 0.0203 | Yes |
| B33 | 36 | 24 | 2.262 | 2.281 | 405.0 | 0.6893 | No |

**Supplementary Table S4. Mann-Whitney U test comparing protein abundance of PhaB** in strain 33B against strains 333, $3B3^B$, $3B3^A$, 4B3, $4B4^A$, and B33, showing sample sizes (N), mean abundances, U statistics, p-values, and significance ($p < 0.05$ indicates a significant difference in distributions).

| Comparator Strain | N (33B) | N (Other) | Mean (33B) | Mean (Other) | U Statistic | p-value | Significance |
|---|---|---|---|---|---|---|---|
| 333 | 27 | 27 | 0.792 | 0.933 | 327.0 | 0.5221 | No |
| $3B3^B$ | 27 | 24 | 0.792 | 0.610 | 459.5 | 0.0108 | Yes |
| $3B3^A$ | 27 | 24 | 0.792 | 0.443 | 543.0 | 0.0000 | Yes |
| $4B3^A$ | 27 | 21 | 0.792 | 0.269 | 543.0 | 0.0000 | Yes |
| 4B4 | 27 | 21 | 0.792 | 0.223 | 565.0 | 0.0000 | Yes |
| B33 | 27 | 18 | 0.792 | 0.805 | 263.0 | 0.6514 | No |

**Supplementary Table S5. Approximate z-test results for log-scale protein ratio of TesB:PhaA** between strain 33B to strains 333, $3B3^B$, $3B3^A$, 4B3, $4B4^A$, and B33, with sample sizes, arithmetic ratios, p-values, and significance ($p < 0.05$ indicates a significant difference in log-ratios).

| Comparator Strain | N_num (33B) | N_den (33B) | N_num (Other) | N_den (Other) | Ratio (33B) | Ratio (Other) | p-value | Significance |
|---|---|---|---|---|---|---|---|---|
| 333 | 27 | 36 | 27 | 36 | 0.845 | 0.626 | 0.5180 | No |
| $3B3^B$ | 27 | 36 | 24 | 32 | 0.845 | 0.698 | 0.4196 | No |
| $3B3^A$ | 27 | 36 | 24 | 32 | 0.845 | 0.522 | 0.2298 | No |
| $4B3^A$ | 27 | 36 | 21 | 28 | 0.845 | 0.250 | 0.0001 | Yes |
| 4B4 | 27 | 36 | 21 | 28 | 0.845 | 0.301 | 0.0014 | Yes |
| B33 | 27 | 36 | 18 | 24 | 0.845 | 0.714 | 0.6986 | No |

**Supplementary Table S6. Approximate z-test results for log-scale protein ratio of TesB:PhaB** between strain 33B to strains 333, 3B3$^{B}$, 3B3$^{A}$, 4B3, 4B4$^{A}$, and B33, with sample sizes, arithmetic ratios, p-values, and significance ($p < 0.05$ indicates a significant difference in log-ratios).

| Comparator Strain | N_num (33B) | N_den (33B) | N_num (Other) | N_den (Other) | Ratio (33B) | Ratio (Other) | p-value | Significance |
|---|---|---|---|---|---|---|---|---|
| 333 | 27 | 36 | 27 | 36 | 0.845 | 0.626 | 0.5180 | No |
| 3B3$^{B}$ | 27 | 36 | 24 | 32 | 0.845 | 0.698 | 0.4196 | No |
| 3B3$^{A}$ | 27 | 36 | 24 | 32 | 0.845 | 0.522 | 0.2298 | No |
| 4B3$^{A}$ | 27 | 36 | 21 | 28 | 0.845 | 0.250 | 0.0001 | Yes |
| 4B4 | 27 | 36 | 21 | 28 | 0.845 | 0.301 | 0.0014 | Yes |
| B33 | 27 | 36 | 18 | 24 | 0.845 | 0.714 | 0.6986 | No |

**Supplementary Table S7. Approximate z-test results for log-scale protein ratio of PhaA:PhaB** between strain 33B to strains 333, 3B3$^{B}$, 3B3$^{A}$, 4B3, 4B4$^{A}$, and B33, with sample sizes, arithmetic ratios, p-values, and significance ($p < 0.05$ indicates a significant difference in log-ratios).

| Comparator Strain | N_num (33B) | N_den (33B) | N_num (Other) | N_den (Other) | Ratio (33B) | Ratio (Other) | p-value | Significance |
|---|---|---|---|---|---|---|---|---|
| 333 | 36 | 27 | 36 | 27 | 2.367 | 3.362 | 0.1929 | No |
| 3B3$^{B}$ | 36 | 27 | 32 | 24 | 2.367 | 3.546 | 0.0802 | No |
| 3B3$^{A}$ | 36 | 27 | 32 | 24 | 2.367 | 3.111 | 0.3459 | No |
| 4B3$^{A}$ | 36 | 27 | 28 | 21 | 2.367 | 3.449 | 0.0648 | No |
| 4B4 | 36 | 27 | 28 | 21 | 2.367 | 3.102 | 0.2341 | No |
| B33 | 36 | 27 | 24 | 18 | 2.367 | 2.832 | 0.4149 | No |

**Supplementary Table S8. Cohen's d effect sizes for protein abundance of TesB**, comparing strain 33B to strains 333, 3B3[B], 3B3[A], 4B3, 4B4[A], and B33, with mean abundances and d values (positive d indicates higher mean in 33B; |d| < 0.2 small, 0.2–0.5 small, 0.5–0.8 medium, > 0.8 large).

| Comparator Strain | Mean (33B) | Mean (Other) | Cohen's d |
|---|---|---|---|
| 333 | 1.942 | 1.649 | 0.205 |
| 3B3[A] | 1.942 | 1.539 | 0.274 |
| 3B3[B] | 1.942 | 0.798 | 0.926 |
| 4B3 | 1.942 | 0.278 | 1.425 |
| 4B4 | 1.942 | 0.230 | 1.468 |
| B33 | 1.942 | 1.629 | 0.216 |

**Supplementary Table S9. Cohen's d effect sizes for protein abundance of PhaA**, comparing strain 33B to strains 333, 3B3$^{B}$, 3B3$^{A}$, 4B3, 4B4$^{A}$, and B33, with mean abundances and d values (positive d indicates higher mean in 33B; |d| < 0.2 small, 0.2–0.5 small, 0.5–0.8 medium, > 0.8 large).

| Comparator Strain | Mean (33B) | Mean (Other) | Cohen's d |
|---|---|---|---|
| 333 | 2.298 | 2.632 | -0.126 |
| 3B3$^{A}$ | 2.298 | 2.204 | 0.040 |
| 3B3$^{B}$ | 2.298 | 1.531 | 0.367 |
| 4B3 | 2.298 | 1.116 | 0.619 |
| 4B4 | 2.298 | 0.766 | 0.835 |
| B33 | 2.298 | 2.281 | 0.007 |

**Supplementary Table S10. Cohen's d effect sizes for protein abundance of PhaB**, comparing strain 33B to strains 333, $3B3^{B}$, $3B3^{A}$, 4B3, $4B4^{A}$, and B33, with mean abundances and d values (positive d indicates higher mean in 33B; $|d| < 0.2$ small, 0.2–0.5 small, 0.5–0.8 medium, > 0.8 large).

| Comparator Strain | **Mean (33B)** | **Mean (Other)** | **Cohen's d** |
|---|---|---|---|
| 333 | 0.971 | 0.783 | 0.430 |
| $3B3^{A}$ | 0.971 | 0.622 | 0.884 |
| $3B3^{B}$ | 0.971 | 0.492 | 1.273 |
| 4B3 | 0.971 | 0.324 | 1.828 |
| 4B4 | 0.971 | 0.247 | 2.107 |
| B33 | 0.971 | 0.805 | 0.364 |

**Supplementary Table S11. Cohen's d effect sizes for log-scale protein ratio of TesB:PhaA** comparing strain 33B to strains 333, 3B3$^{B}$, 3B3$^{A}$, 4B3, 4B4$^{A}$, and B33, showing arithmetic ratios and d values (positive d indicates higher ratio in 33B; |d| < 0.2 small, 0.2–0.5 small, 0.5–0.8 medium, > 0.8 large).

| Comparator Strain | Ratio (33B) | Ratio (Other) | Cohen's d |
|---|---|---|---|
| 333 | 0.845 | 0.626 | 0.521 |
| 3B3$^{B}$ | 0.845 | 0.698 | 0.568 |
| 3B3$^{A}$ | 0.845 | 0.522 | 0.657 |
| 4B3$^{A}$ | 0.845 | 0.250 | 1.401 |
| 4B4 | 0.845 | 0.301 | 1.218 |
| B33 | 0.845 | 0.714 | 0.450 |

**Supplementary Table S12. Cohen's d effect sizes for log-scale protein ratio of TesB:PhaB** comparing strain 33B to strains 333, $3B3^B$, $3B3^A$, 4B3, $4B4^A$, and B33, showing arithmetic ratios and d values (positive d indicates higher ratio in 33B; |d| < 0.2 small, 0.2–0.5 small, 0.5–0.8 medium, > 0.8 large).

| Comparator Strain | Ratio (33B) | Ratio (Other) | Cohen's d |
|---|---|---|---|
| 333 | 2.000 | 2.105 | -0.556 |
| $3B3^B$ | 2.000 | 2.477 | -0.583 |
| $3B3^A$ | 2.000 | 1.623 | -0.285 |
| $4B3^A$ | 2.000 | 0.861 | 0.240 |
| 4B4 | 2.000 | 0.933 | 0.246 |
| B33 | 2.000 | 2.023 | -0.498 |

**Supplementary Table S13. Cohen's d effect sizes for log-scale protein ratio of PhaA:PhaB** comparing strain 33B to strains 333, $3B3^B$, $3B3^A$, 4B3, $4B4^A$, and B33, showing arithmetic ratios and d values (positive d indicates higher ratio in 33B; |d| < 0.2 small, 0.2–0.5 small, 0.5–0.8 medium, > 0.8 large).

| Comparator Strain | Ratio (33B) | Ratio (Other) | Cohen's d |
|---|---|---|---|
| 333 | 2.367 | 3.362 | -0.332 |
| $3B3^B$ | 2.367 | 3.546 | -0.459 |
| $3B3^A$ | 2.367 | 3.111 | -0.219 |
| $4B3^A$ | 2.367 | 3.449 | -0.456 |
| 4B4 | 2.367 | 3.102 | -0.287 |
| B33 | 2.367 | 2.832 | -0.204 |
